# Default Mode Network Connectivity Predicts Individual Differences in Long-Term Forgetting: Evidence for Storage Degradation, not Retrieval Failure

**DOI:** 10.1101/2021.08.04.455133

**Authors:** Yinan Xu, Chantel Prat, Florian Sense, Hedderik van Rijn, Andrea Stocco

## Abstract

Despite the importance of memories in everyday life and the progress made in understanding how they are encoded and retrieved, the neural processes by which declarative memories are maintained or forgotten remain elusive. Part of the problem is that it is empirically difficult to measure the rate at which memories fade, even between repeated presentations of the source of the memory. Without such a ground-truth measure, it is hard to identify the corresponding neural correlates. This study addresses this problem by comparing individual patterns of functional connectivity against behavioral differences in forgetting speed derived from computational phenotyping. Specifically, the individual-specific values of the speed of forgetting in long-term memory (LTM) were estimated for 33 participants using a formal model fit to accuracy and response time data from an adaptive paired-associate learning task. Individual speeds of forgetting were then used to examine participant-specific patterns of resting-state fMRI connectivity, using machine learning techniques to identify the most predictive and generalizable features. Our results show that individual speeds of forgetting are associated with resting-state connectivity within the default mode network (DMN) as well as between the DMN and cortical sensory areas. Cross-validation showed that individual speeds of forgetting were predicted with high accuracy (*r* = .78) from these connectivity patterns alone. These results support the view that DMN activity and the associated sensory regions are actively involved in maintaining memories and preventing their decline, a view that can be seen as evidence for the hypothesis that forgetting is a result of storage degradation, rather than of retrieval failure.

**Author Summary:** Why do some people forget faster than others? This study investigates individual differences in long-term memory forgetting by linking them to patterns of brain connectivity. Although memory formation and retrieval are well-studied, much less is known about the brain processes that determine how memories are maintained or lost over time. A major challenge has been accurately measuring forgetting. To address this, we used an innovative method: fitting a computational model of memory to each participant’s accuracy and response time data during a learning task. This allowed us to estimate each person’s unique rate of forgetting. We then examined how these rates related to resting-state brain connectivity. The results revealed that individual forgetting speeds were strongly associated with connectivity between the brain’s default mode network (DMN) and cortical sensory regions—areas thought to support long-term memory maintenance. These findings suggest that forgetting reflects how well memory traces are preserved in the brain, not just whether they can be retrieved.

## Introduction

Despite considerable progress in memory research, we still struggle to understand how and why declarative memories (that is, semantic and episodic memories [1]) are lost and *forgotten*. In fact, researchers disagree even about when and how forgetting takes place.

Consider, for example, the three-stage model of a memory’s lifecycle, which is a staple of cognitive psychology textbooks [1]. In this idealized depiction, memories are first *encoded* within newly formed synapses that connect a network of different neurons. This neural representation is then *stored* and maintained for an extended amount of time, during which their neural representations might undergo significant changes. Finally, memories are *retrieved* at some later time—an interval which can vary from minutes to years. Because forgetting is only inferred after a failed retrieval attempt, it is difficult to pinpoint at which of these stages it really occurred. Specifically, researchers disagree on whether forgetting is a process that is best thought of as degradation occurring during the storage stage [2,3] or as a consequence of failures in the retrieval stage [4–6] .

According to the *storage degradation* class of theories, forgetting mostly interests how memories are maintained during the storage phase. In this view, the neural representations of memories are intrinsically labile and accumulate progressive damage over time. Such damage could be due, for example, to biological decay processes, such as the spontaneous loss of synapses [3,7,8]. It could also be due to cells and synapses of the original neural representation being recruited to encode new traces (a form of catastrophic interference: [9]). Finally, memories might be damaged during *system consolidation* , a post-encoding process that alters the underlying representations of a memory, resulting in a re-organization of the cell assemblies involved. Spontaneous brain activity at rest, for example, is believed to play a pivotal role in system consolidation [10], and its disruption is associated with abnormal forgetting in amnestic neurodegeneration [11,12].

According to the *retrieval failure* class of theories, the underlying neural representations of memories are relatively stable over time, and forgetting is instead driven by an inability to properly access and retrieve them [13]. As in the case of storage degradation, multiple processes have been hypothesized to explain retrieval failure. For example, memories might become less accessible because of interference, that is, the accumulation of competing memories that hinder the retrieval process. Most theories assume controlled memory retrieval is underpinned by the prefrontal cortex; thus, it is possible that loss of synaptic connections between prefrontal regions and the cell assemblies where memories are stored renders them inaccessible. Evidence for these theories comes from the fact that direct neurostimulation of sensory regions might sometimes re-awaken long-forgotten memories, a fact that has been reported in humans [14,15] (although not without criticism: [16,17]) and, to the extent to which evidence for episodic memory can be ascertained in animals, in rodent models of Alzheimer’s Disease [18].

### Implications and Predictions of Storage Degradation vs. Retrieval Failure

The distinction between these two families of theories is not merely academic; it has important implications for both memory research and clinical practice. Consider, for example, Alzheimer’s Disease (AD), a dramatic form of abnormal forgetting. If the storage degradation theory is correct, the promising new clinical treatments that remove amyloid beta deposits (e.g., [19,20]) would stop the progression of the disease but would not restore memory function and would not bring back lost memories, as the underlying neural substrate has already been compromised. However, according to the retrieval failure theory, at least some lost memories might be retrievable, and function could be partially restored. Knowing which theory is correct, therefore, is important to guide treatment and future clinical research.

The current understanding of the mechanisms by which memories are encoded, maintained, and retrieved makes it possible to distinguish between these two theories. Most theories assume that the initial representation of a memory occurs in the hippocampus, by creating synapses between hippocampal cells that are also connected to the sensory areas that were initially involved at the moment of encoding [21,22]. The re-activation of this cell assembly, sometimes referred to as *engram* [4,8,23,24], results in the re-experience of the associated memory [25,26]. Replays of this activity, either through active retrieval or spontaneous brain activity, result in consolidation. This internal re-activation of the memories is supported by the coordinated activity of the so-called Default Mode Network (DMN), a large-scale brain network that includes the hippocampus, the medial frontal cortex, and the precuneus [27]. The DMN is believed to play a key role in propagating the re-activation of the original engrams, mediating the interactions between the hippocampus and the neocortical regions during consolidation [28]. In fact, the DMN’s activity at rest is explained as a way of rehearsing and maintaining memory as a form of “priors” for later processing [10,29,30].

The controlled retrieval of declarative memories, on the other hand, is a multifaceted process supported by the activity of the DMN and multiple dorsal and lateral prefrontal circuits. The DMN might be involved in mediating access to declarative memories, and episodic traces in particular [31–33]. Dorsal prefrontal regions are involved in voluntary and controlled memory retrieval as well as working memory, and provide a natural connection between the two forms of memory [34]. Ventral lateral prefrontal regions, on the other hand, are specifically involved in the selection between competing memories [35], including the resolving conflict between potential candidates [36] and maintaining retrieval cues [37–39]. Ventral and medial prefrontal regions, finally, seem to play a critical role in monitoring the retrieval process, as evidenced by their involvement in meta-memory judgments and feeling-of-knowing [40–43] and in confabulation in frontal patients [44,45]. Damage to these frontal regions, as well as the degradation of connectivity between these regions and the DMN, might result in the appearance of forgetting even when the underlying neural representations are intact—and might be directly awakened through electrical stimulation [14].

Since consolidation and retrieval processes have different neural substrates, the storage degradation and retrieval failure theories also make different predictions in terms of the neural correlates of forgetting. According to the storage degradation framework, individual differences in forgetting should be accompanied by differences in the functional connectivity between the DMN [28,46] and the sensory and association areas where the individual features of a memory are initially encoded—and where long-term memories might be stored as the end-product of systems consolidation [47–49].

According to the retrieval failure framework, however, forgetting occurs because of an inability to voluntarily access previously stored memories, even if the same memories can be involuntarily re-activated (for example, through local brain stimulation). Because the controlled retrieval of memories is controlled by lateral prefrontal cortices [34,35,39], the theory predicts that forgetting would be associated with the functional connectivity between the prefrontal and parietal regions that oversee controlled retrieval and the encoding and consolidation regions of the DMN [50].

### Obstacles to the Testing the Theories

Since the two theories make different predictions, it should be possible to empirically test them by examining which features of individual anatomy and functional connectivity co-vary with individual differences in forgetting. One important obstacle, however, remains: it is extremely difficult to *quantify* forgetting in the first place. As noted above, forgetting is typically operationalized as failing to correctly recall or recognize an item in a memory test: performance failure is the only ground truth available. Unfortunately, this carries significant limitations. First, common behavioral measures such as the proportion of correctly remembered items are prone to statistical misinterpretations [51,52]. Second, performance in memory tests is also affected by the specifics of the paradigms used. For example, memory performance differs greatly when the same materials are tested using recognition or recall; recognition is generally easier, but recall, being more effortful, leads to better subsequent consolidation and retention.

This retrieval accuracy-based operationalization does not easily account for possible temporal dynamics of forgetting. If forgetting is conceptualized as a process, a more precise assessment of memory accessibility is needed even if a memory test results in a successful retrieval.

The goal of this study is to overcome these limitations through *computational phenotyping* [53,54], that is, by fitting a formal model of cognitive processes to individual task data, and using the model’s parameters to quantify individual differences. Model parameters obtained through computational phenotyping are easier to interpret and more reliable than raw behavioral measures [54–56], such as accuracies or response times. In this study, individual differences in forgetting were quantified by fitting a computational model of forgetting to individual learning data in a sample of healthy individuals for whom functional connectivity data

were available. By analyzing which connectivity features best predict individual differences in forgetting, we can empirically compare the storage degradation and retrieval failure theories.

### A Computational Theory of Forgetting

As referred to above in the context of proportion of correct responses, purely behavioral measures of psychological constructs often have poor reliability [52], which leads, in turn, to poor brain-behavioral associations [57]. One way to obviate this limitation is to fit realistic computational models to individual behaviors, and use the fitted model parameters as dependent variables [58,59]. When the model captures the dynamics of the underlying neural process, its parameters can be more reliable and interpretable than pure behavioral measures. This has been shown, for example, with the threshold and drift parameters of accumulator models in perceptual decision-making [60,61] and in learning rate parameters in reinforcement learning models of learning [56]. Finally, using realistic models of long-term memory avoids the pitfalls that were originally pointed out by Loftus [51,62] in applying statistical linear models to non-linear effects in memory performance.

In this study, the estimation of the speed of forgetting was based on a well-established model of episodic memory [55,63–66] that is also the cornerstone of the ACT-R theory [67–69]. This model, coupled with an adaptive learning task that dynamically adjusts the presentation schedules of the stimuli [70], yields individual speeds of forgetting for individuals that can be measured within a short time (< 15 mins) but can be reliably extrapolated over much longer intervals spanning days or weeks [55,66,71,72].

In the model, the probability of retrieving a memory *m* at time *t* is proportional to its activation *A*(*m, t*), a scalar quantity that reflects a memory’s availability. As in the multiple trace theory [73,74], each memory *m* is associated with multiple episodic *traces*, each of which corresponds to specific (but potentially overlapping) neural representations that were created when *m* was retrieved or (re)encoded, and each of which decays over time.

The activation of each trace decays as a power function of time [75], and the activation of *m* is the log sum of the decaying activations of all its traces:

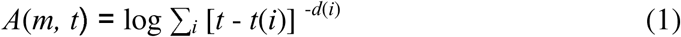

where *t*(*i*) and *d*(*i*) are, respectively, the time of the creation of the *i*-the trace and its characteristic “decay rate”. The decay rate can be thought of as the degree of damage (caused by either biological decay, system consolidation, or interference) that a trace accumulates per unit time, and which reduces that trace’s quality and, therefore, retrievability [69]. A trace’s decay rate, in turn, is related to the *m*’s activation at the moment each trace is created, thus capturing the fact that traces with higher initial activation decay faster than traces with lower activation [64]. Specifically:

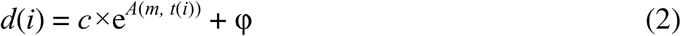

with *A*(*m*, *t*(*i*)) being the activation of *m* at the moment the *i*-th trace was encoded, and *c* is a scaling factor fixed to 0.25, as previously proposed by [70]. Thus, if the times at which a memory *m* has been encoded, re-encoded, and retrieved are known and controlled for, its activation depends only on φ, which is the only free parameter in the above equations. After determining the φ of each memory, their average was taken to represent an individual-specific *speed of forgetting* (SoF).

The activation of a memory, thus defined, determines its retrieval probability. Specifically, the retrieval probability *P*(*m*) of a memory *m* can be approximated as:

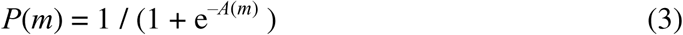

Previous studies have shown that individual values of φ are stable across time and materials and predictive of real-world outcomes such as learning and test scores [55,66,76–78], thus meeting the psychometric criteria of reliability and validity [79]. Finally, at least one previous study has established that speed of forgetting, thus measured, reliably correlates with individual differences in EEG activity, thus establishing a connection between SoF and brain function [80].

### Experimental Predictions

To distinguish between the retrieval failure and storage degradation hypotheses, this study applied a statistical learning approach [81] to resting-state fMRI data from healthy participants. Specifically, a regularized linear model’s hyperparameter was tuned to minimize the cross-validated prediction error of an individual’s speed of forgetting from their resting-state connectivity. The specific patterns of connectivity that were identified to successfully predict individual differences in forgetting would shed light onto the biological nature of the phenomenon by indicating whether the involved networks align best with the retrieval or the storage degradation hypothesis.

Resting-state activity fMRI was chosen because of its spatial precision and its reliability within participants [82]. It has been speculated that brain activity at rest reflects the invariant functional architecture of the brain [30,83]; small variations within this architecture are characteristic to every individual, to the point that it can be used to “fingerprint” individual participants [84,85]. Consistent with this finding, individual differences in the resting-state functional connectivity between different regions have been shown to be predictive of performance differences in a variety of cognitive abilities, including working memory [86], executive functions [87,88], motor learning [89], perceptual discrimination [90], and, most importantly, recognition memory [91]. Thus, resting-state functional connectivity provides an ideal target to identify the biological underpinnings of individual differences in long-term memory retention and forgetting.

Specifically, we hypothesized that the speed of forgetting would reliably co-vary with the functional connectivity between different sub-networks of the DMN. This prediction is common to both hypotheses and follows from the central role played by the DMN in systems consolidation and episodic memory retrieval.

In addition, we also hypothesized that individual differences in the speed of forgetting will be reflected in the connectivity between the DMN and other networks. These patterns of connectivity would allow us to distinguish between the storage degradation and the retrieval failure hypotheses. If the storage degradation framework is correct, then we expect to see that individual differences in forgetting are best predicted by patterns of connectivity between the DMN, sensory, and associative cortical areas [49,74,92], including higher-level visual areas such as those associated with the ventral and dorsal visual attention networks.

If, on the other hand, the retrieval failure framework is correct, then we would expect to find that individual differences in forgetting are best predicted by the connectivity between the DMN and brain networks associated with frontal and parietal control regions.

## Materials and Methods

### Participants

A total of 33 monolingual English-speaking participants (19 females) aged between 18 to 33 years old were recruited from a pool of University of Washington undergraduates for whom resting-state fMRI data had already been acquired in a previous, unrelated experiment between 6 and 12 months before the paired-associate learning task was administered. All participants provided informed consent and were compensated with a $25 gift card for their participation in the paired-associate study. All of the recruitment and testing procedures were approved by the University of Washington’s Institutional Review Board.

### Adaptive Paired Associate Learning Task

Individual speeds of forgetting were estimated using the adaptive paired-associate learning task described in [55]. The stimuli consisted of 25 Swahili-English word pairs selected from a previous study [93,94]. The task dynamically interleaved study trials and test trials following a retrieval practice design [95]. On study trials, a Swahili word (e.g., “samaki”) and its corresponding English equivalent (“fish”) were presented simultaneously on the screen, and participants typed in the English word to proceed. On test trials, only the Swahili word was presented on the screen as a cue (e.g., “samaki”), and participants were again asked to respond by typing the corresponding English word (“fish”) in an empty textbox. Trials were self-paced, with no cap on the time allowed for a response. Consecutive trials are separated by a 600 ms ISI after a correct response and a participant paced ISI following an incorrect response during which corrective feedback was shown on screen. The order of repetitions and moment of introduction for each item were determined by the adaptive scheduling algorithm outlined below and described in more detail elsewhere [71,76]. The task lasted 12 minutes, during which participants learned as many of the paired associates as possible. In the allotted time, participants learned, on average, 23.3 different pairs (range: 10-25), across 142.7 trials (range: 91-221), with a mean accuracy of 86.5% on test trials (range: 62.4% - 98.9%).

The scheduling algorithm was designed to optimally interleave the stimuli so that items would be repeated before the internal cognitive model predicted that they would be forgotten as determined by the continuously updated speed of forgetting (φ) for each item. In the model, each memory *m* is a paired associate (see [96]) that links a new Swahili word (“samaki”) to a known English word (“fish”). In response to a study probe, participants are given the Swahili word (“samaki”) as a cue to retrieve *m* (“samaki” - “fish”). Only when *m* is retrieved can participants type in the correct answer (“fish”). In turn, φ is estimated from the time *T* it takes participants to respond (first key press), which is related to each pair’s activation by the equation:

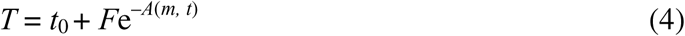

where *A*(*m, t*) is the current memory activation (Eq. 1), *F* is a scaling constant fixed to 1, and *t*_0_ represents a fixed offset (300 ms) for non-memory related processes, such as the time spent for encoding the visual stimulus and making a motor response [68,97]. Thus, responses that are either incorrect or slower than expected result in upwards adjustments of φ, while responses that are faster than expected result in downwards adjustments of the parameters.

Although it might seem surprising that long-term memory forgetting can be assessed within such a relatively short period, this is a consequence of the fact that forgetting follows a power law [75], in which forgetting occurs most quickly in the first few minutes and then gradually slows down. For example, in a forgetting curve with an SoF value of 0.3 (which is a reasonable estimate for the average φ for undergraduate students; in this study the average is 0.305, see below), a memory’s retrieval probability falls < 25% within the first 10 mins, unless it is rehearsed. This means that the first minutes are an ideal interval to precisely estimate the speed of forgetting, as a significant part of the decline happens in this short timeframe.

### rs-fMRI Data Collection and Preprocessing

Each resting-state run lasted 7 minutes and consisted of 210 echo-planar images (EPI) collected with TR = 2,000 ms, TE = 25 ms, FA = 72 degrees. Each image in the series consisted of 36 oblique 3-mm bi-dimensional slices with no gap in-between, with an in-plane resolution of 80x80 3x3 mm voxels. Each functional series was preprocessed using the AFNI [98] software package. Specifically, each resting-state time series was de-spiked, corrected for differences in slice acquisition time, realigned to the mean image in the series, normalized to the MNI ICBM152 template, and spatially smoothed with a 3D 8-mm FWHM Gaussian filter. Artifacts induced by subject movement were removed by regressing out the six motion parameters and their temporal derivatives from each voxel’s time series. In addition to the functional data, a high-resolution anatomical MP-RAGE was also collected to aid in the normalization process.

### Brain Parcellation

To calculate functional connectivity, each participant’s brain was divided into discrete regions using the network parcellation originally proposed by Yeo et al [99]. Figure 3 depicts the 17 brain networks identified by Yeo, using the color scheme proposed in the original paper.

**Figure 1:**
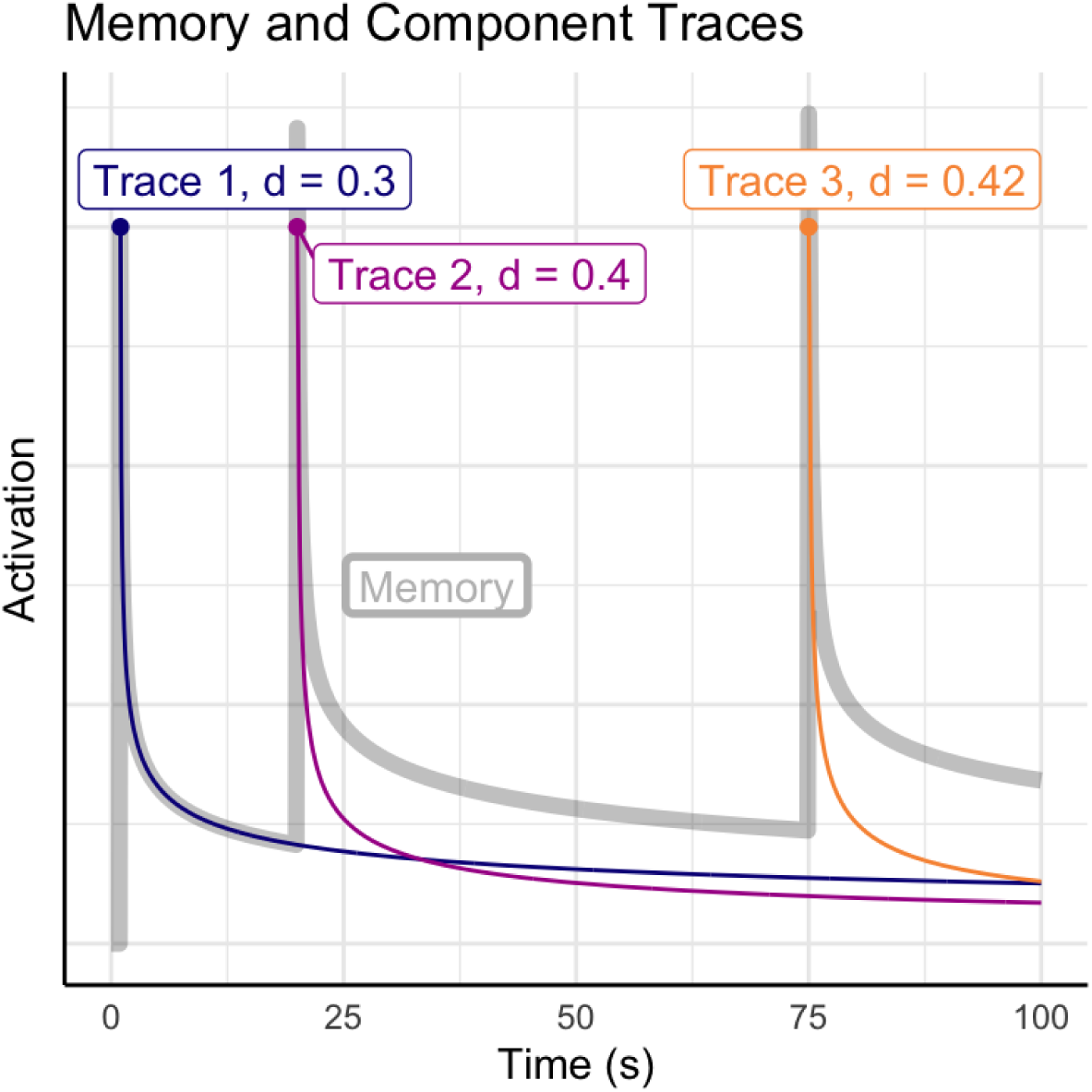
Activation over time of a memory that has been encoded in three different episodes at times t = 1, 20, and 75 seconds. Each episode leaves an independent memory trace (colored lines) with a characteristic decay rate. The total activation of the memory is the sum of the decaying traces (gray line).

**Figure 2.**
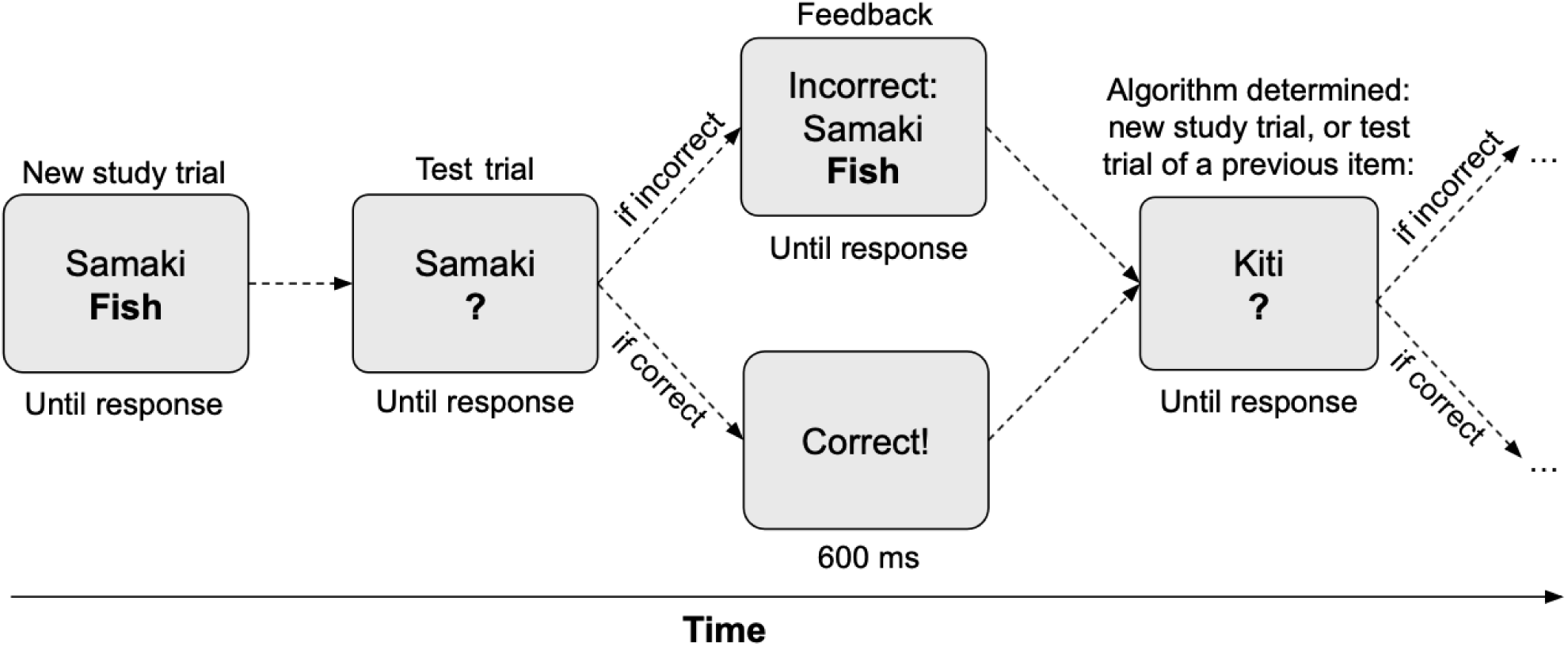
Visual illustration of one complete study trial and the onset of a test trial from the adaptive vocabulary learning task. Note that study items are repeated based on an estimate of when their memory activation would drop below a predefined threshold.

**Figure 3.**
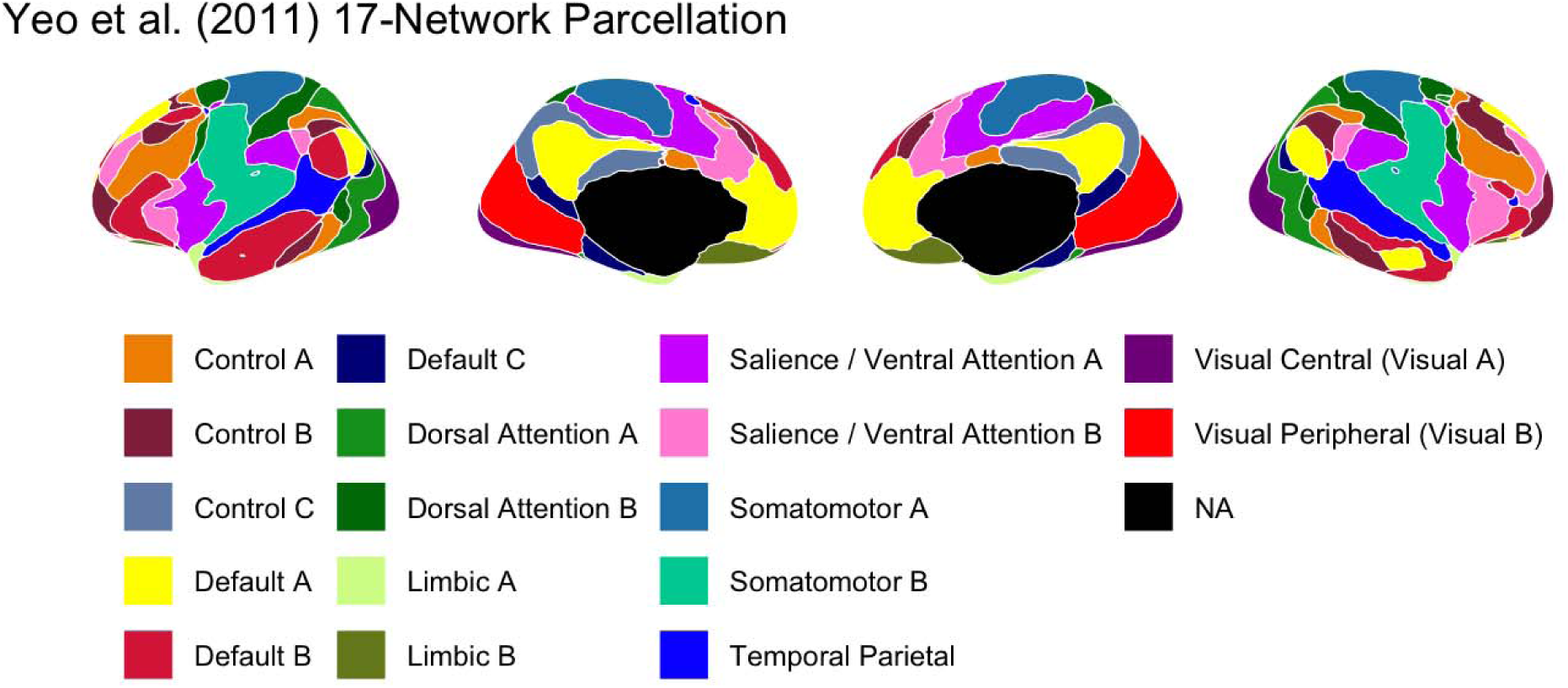
The 17 networks in the Yeo et al (2011) parcellation. Subcortical regions, not included in the parcellation, are marked in black.

To test the storage degradation and retrieval failure hypotheses, each network was assigned to one of three possible groups:

● The Default Mode Network, consisting of Default A, B, and C as identified by [99];
● The Retrieval group, consisting of networks including key prefrontal and parietal regions implicated in controlled and voluntary memory retrieval [34,35,37,39,50] as well as the ventral and medial prefrontal regions monitoring of the retrieval process [41–43]. These regions include the subnetworks Control A and B, Dorsal Attention A, Salience B, Limbic B, and Temporal Parietal in the Yeo parcellation.
● The Storage group, consisting of regions associated with the encoding of the memory’s sensory and affective features and its later consolidation of memory; thus, they encompass both perceptual and sensorimotor areas [100–106] and the insular and medial regions involved in assessing stimulus salience [107–109]. These regions include Yeo’s subnetworks Visual A and B, Somatomotor A and B, Dorsal Attention B, Salience A, Limbic A, and Control C.

As would be expected of functionally related regions, the retrieval and storage networks, thus defined, cluster into spatially contiguous portions of the brain, as illustrated in Figure 4.

**Figure 4:**
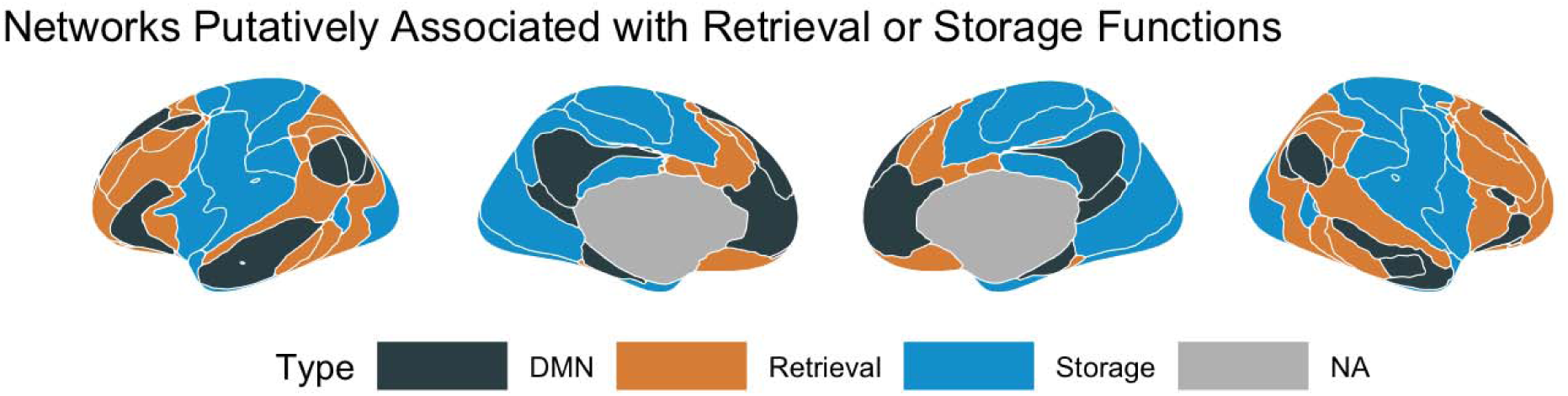
The grouping of the networks from the Yeo et al (2011) parcellation according to their involvement in consolidation (DMN, black) storage (blue) or retrieval (brown) processes.

**Figure 5:**
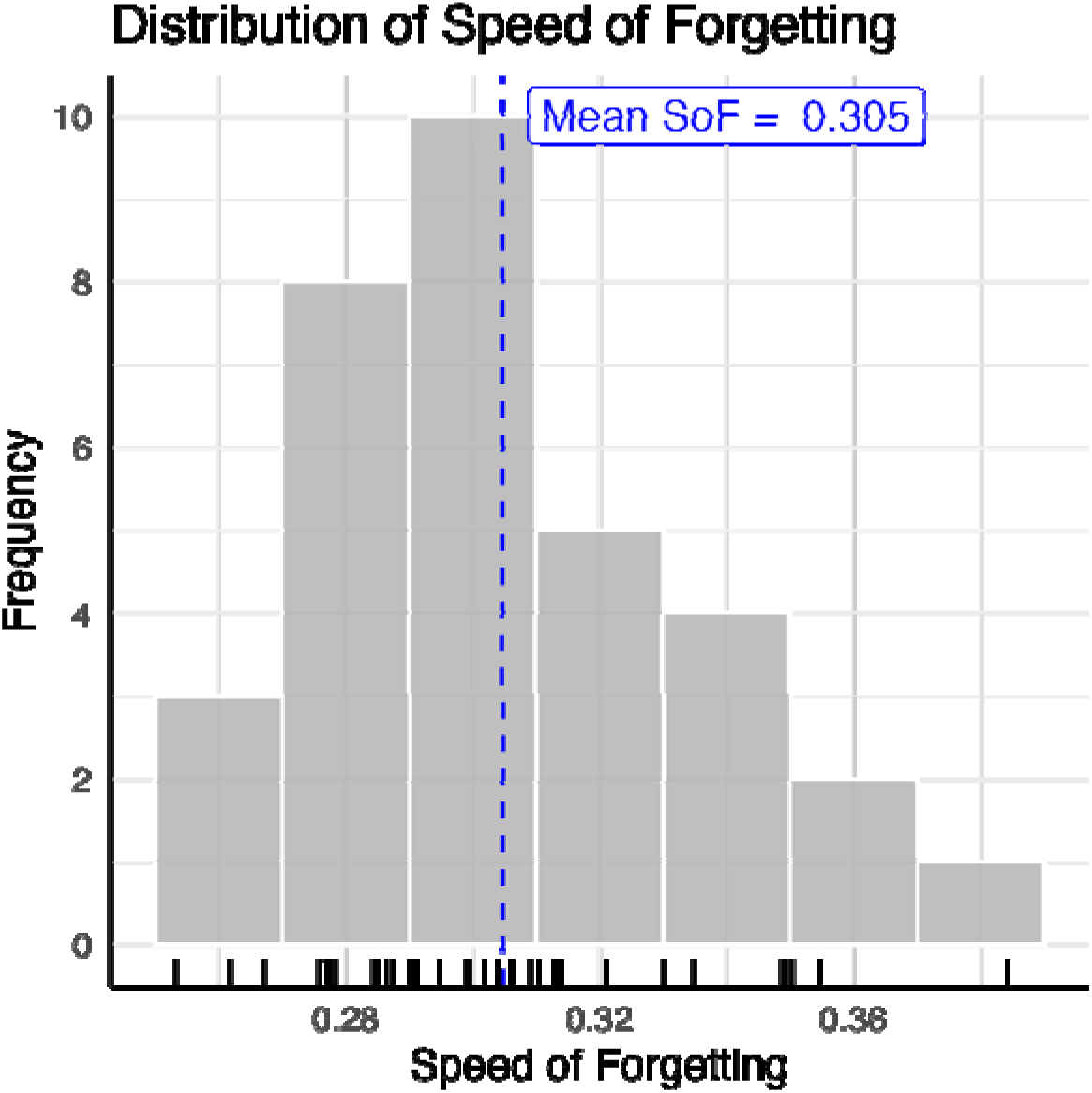
Distribution of the observed speeds of forgetting (N=33). Rug plot lines on the x-axis indicate values from each individual participant.

### Connectome Extraction

Each participant’s global pattern of functional connectivity, known as the *connectome* [110], was extracted using a standard procedure [111,112]. For each of the 17x17 pairs of regions in the parcellation, the Pearson correlation coefficient between the respective time series was computed.

One problem with this canonical connectome is that the raw correlation coefficients are almost all positive and partially driven by common, correlated factors. To remove these confounds, we followed the procedure of Cole and colleagues [111], and recalculated the correlation matrices using *partial* correlation coefficients, so that, when computing the correlation between two networks, the correlations between each network in the pair and the remaining 15 networks were partialled out. The partial correlation coefficients were the functional connectivity data that was fed into the statistical analysis (see below).

### Statistical Analysis

Functional connectivity data were analyzed with the Least Absolute Shrinkage and Selection Operator [LASSO: 113], a statistical learning method [114,115] that combines linear regression and feature selection in a single framework. Like linear regression, LASSO estimates a vector of regression weights β**\*** that minimizes the squared sum of the differences between the observations **y** and their predicted values, that is argmin_β_||**y** – β**X**||_2_ (where the notation ||v||_n_ represents the L*n* norm of the vector **v**). Unlike traditional regression, the difference is further penalized in proportion to the sum of the absolute values of the regressor weights (their L1 norm), so that:

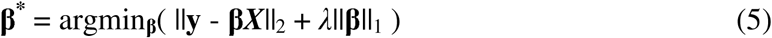

The use of the L1 norm allows the least informative regressors to shrink to zero and, effectively, be dropped from the model—thus combining shrinkage and feature selection [113].

In turn, this reduces the complexity of the model, favoring generalizability and reducing overfitting. The hyperparameter λ represents the tradeoff between model simplicity (encouraged by the ||β||_1_ penalty) and accuracy (encouraged by the least-squares minimization term ||**y** – β**X**||_2_). As with all machine learning approaches, careful validation is needed to estimate the values of λ and to identify the proper predictors; in this project, Leave-One-Out cross-validation was used to select the proper value of λ by minimizing the prediction error over the out-of-sample observation; the result of this process, therefore, is a set of sparse regressors that have the greatest predictive weight. Once the best value of λ was identified, a final model was then constructed by refitting the entire dataset.

## Results

### Speed of Forgetting

One SoF parameter was calculated for each individual by averaging across the φ values of each word pair they studied and which was presented at least three times. The resulting values ranged from 0.253 to 0.385, with a mean of 0.305 and a standard deviation of 0.029. The mean and the range of values are in line with previously published studies [55,80]; no difference was found between male and female participants (Male: 0.309 ± 0.029); Female: 0.304 ± 0.031; *t*(24.47) = –0.48, *p* > 0.63).

It is worth examining how well the model, and especially its predicted SoF parameters, actually fit the participants’ data. Because they responded correctly in > 86% of the trials (close to ceiling due to the adaptive nature of the task), we focused on the model’s predicted response times. To do so, we recalculated the predicted activation for each test trial of every participant, using Eq. 1 and each participant’s final SoF^1^. Then, for each of the 3,354 trials, we computed the predicted response times using Eq. 4 and keeping its remaining parameters to the same default values (*t*_0_ = 300 ms and *F* = 1) used during the task. Response times taking longer than 8 seconds were excluded as outliers; resulting in the removal of 155 trials (4.4% of the total). As it is common for computational models of behavior, we estimated the model’s fit by comparing the predicted and observed distributions of response times; in this case, we used the Kullback-Lieber (KL) divergence metric and the χ^2^test [116]. KL values can be interpreted, in information theory, as the mean number of bits lost when the observed response times are replaced by the model’s predictions, while the χ^2^ values quantify how likely it is that the observed and predicted values are generated by the distribution; in both cases, values closer to zero are better. To compute both metrics, predicted and observed response times were divided into ten 800ms bins, and the number of predicted vs. observed values falling into each bin was compared. As shown in Figure 6, the model is successful at capturing individual differences in response times, producing remarkably matching distributions for each participant. The mean KL value was 1.61 across all participants and 0.35 across all trials from all participants. Similarly, the mean χ^2^ value was 25.4 across participants and 42.0 across all trials, implying that the predicted and observed distributions of response times were not significantly different (that is, *p* > 0.05). Taken together, the results of this analysis confirms not only the validity of the model from which the SoF values were derived, but also the choice of the default parameters used in the adaptive paired associate learning paradigm.

**Figure 6:**
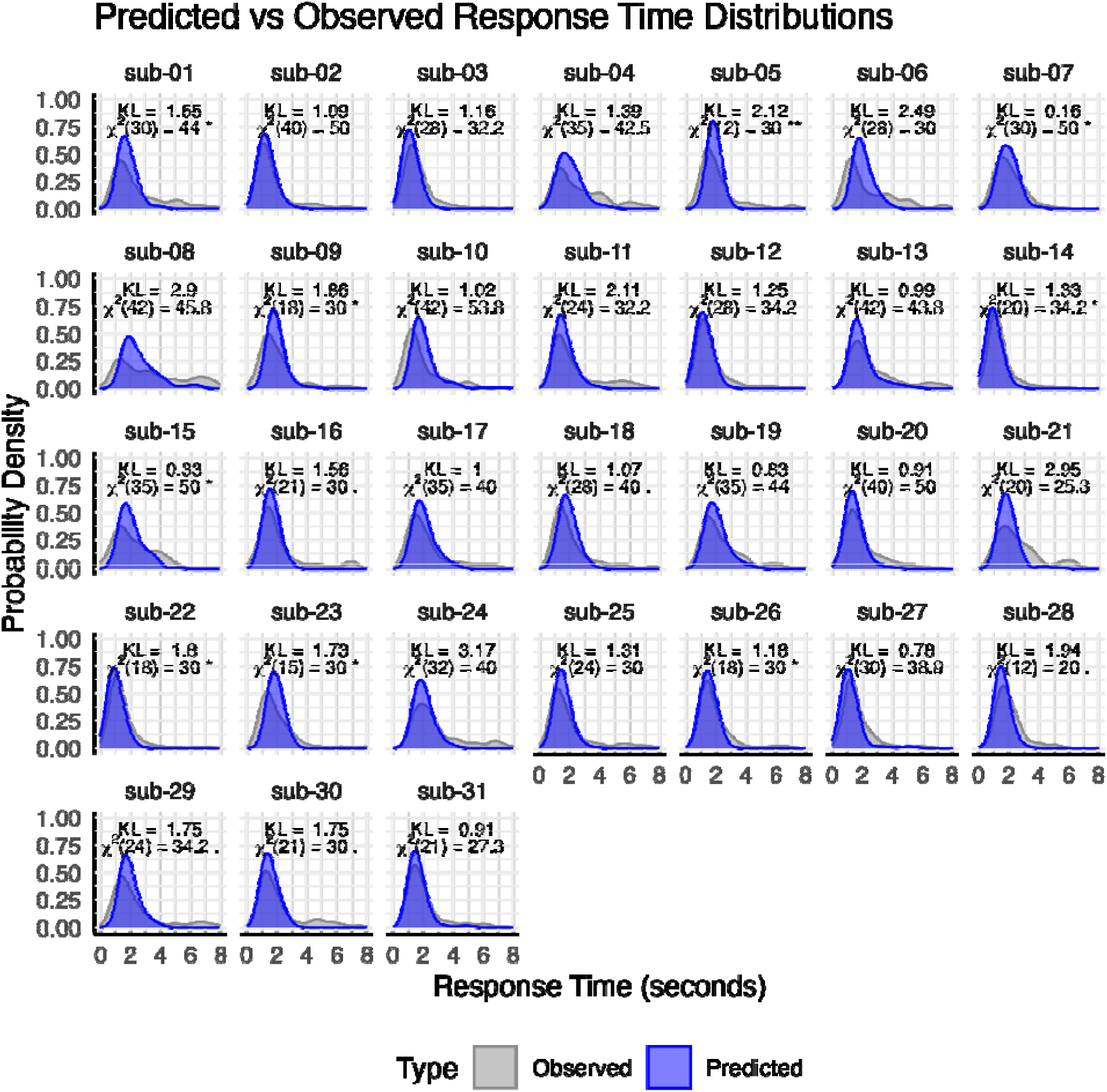
Predicted vs. Observed distributions of response times in the adaptive paired associate learning task. The predicted values are derived from the memory model (Equations 1–4), using participant-specific values for the SoF parameter.

### Resulting Connectome and LASSO Model

Figure 7 shows the group-averages of the resulting individual connectome matrices. The left side shows a canonical distribution of *r* values, including the zones of maximum *r* values along the diagonal (corresponding to connectivity within the same networks). This pattern is entirely consistent with previous findings using the same parcellation scheme.

**Figure 7.**
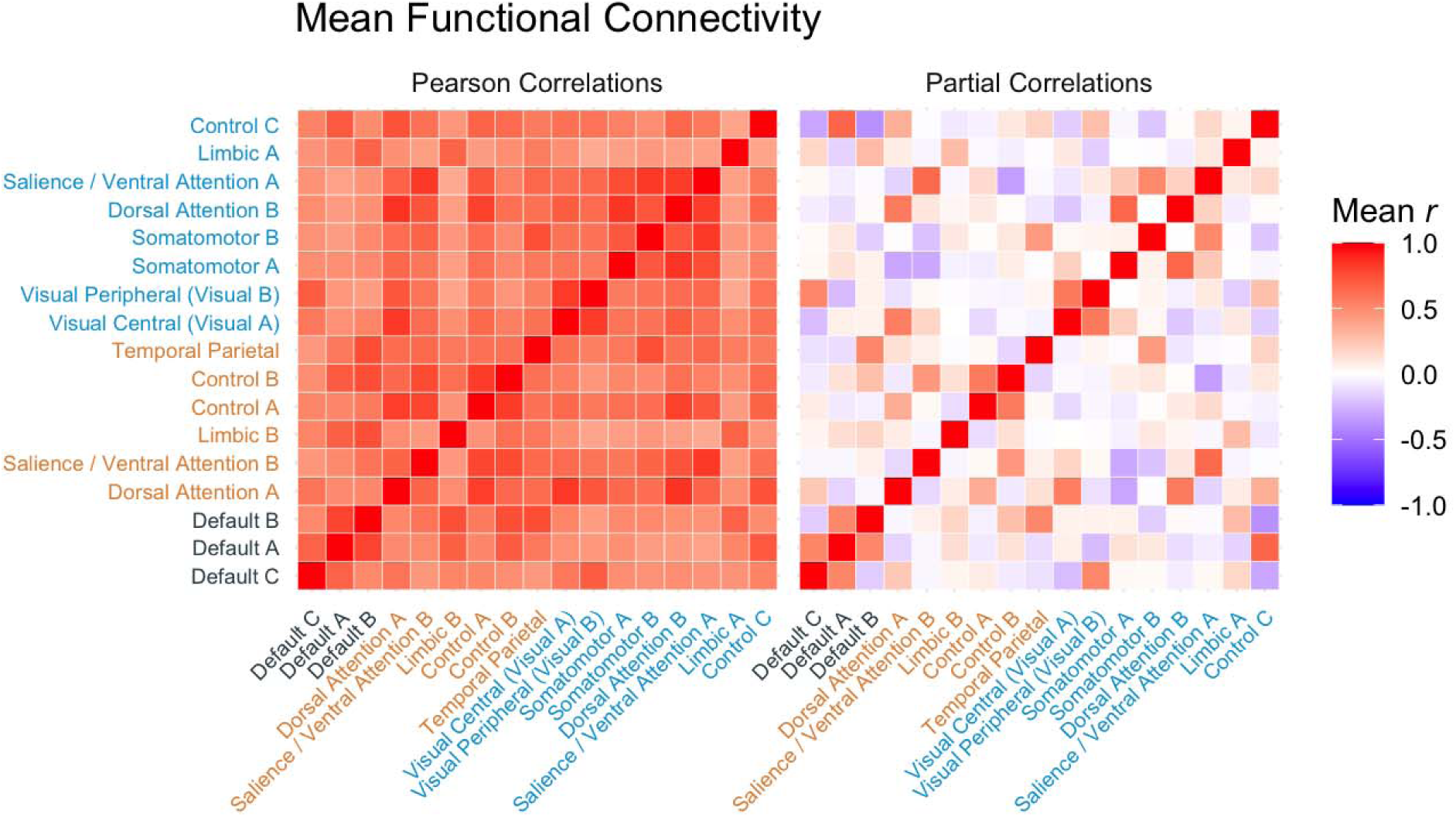
Mean raw Pearson correlations (left) and partial correlations (right) between each of the 17 networks across all participants. The mean correlation coefficient across participants was calculated by first turning each individual coefficient into a z-value using Fisher’s r-to-z transformation, then averaging the individual z-values, and finally transforming the average z value back into an equivalent r coefficient. Network names are color-coded based on whether the network belongs to the Storage, Retrieval, or DMN group (Figure 4).

The right side of Figure 7 shows the average of the partial correlation connectomes. As expected, this second matrix is sparser and includes both negative and positive correlations [111,117], giving a clearer picture of the underlying functional organization. The raw partial connectomes were the features entered into the LASSO model.

Although LASSO reduces the number of predictors, it has been shown that its performance is hindered when the number of regressors is much larger than the number of participants [118], which would be the case if we considered the full 17 x 16 / 2 = 136 possible connections between networks. Thus, to reduce the number of regressors, we limited the analysis to the connections between the three default mode networks (Default A, B, and C in Figure 1), including their connectivity with each other, and all of the other 14 networks. Thus, the total number of regressors included in this analysis was 3 x 14 + 3 = 45. Notice that this does not reduce the generality of our findings, since our theoretical hypotheses concern precisely the relationship between the default mode networks and other networks (associative vs. control networks).

### Network Connectivity Predictive of Individual Speed of Forgetting

The cross-validation procedure yielded a hyperparameter value of λ = 0.0044, which resulted in only 8 regressors representing network connections being selected: it was the individual variations in these regressors that were found to be most predictive of individual speeds of forgetting. Table 1 lists these eight network connections, while Figure 7 provides a visual breakdown of their connectivity, organized according to the different DMN subnetworks involved. All of the six networks whose connectivity to the DMN was identified as a reliable predictor of forgetting (Somatomotor A, Visual A, Salience A, Control C, Limbic A, and Dorsal Attention B: Table 1 and Figure 8) belong to the storage-related group (see Figure 4), suggesting that forgetting can be better understood within the storage degradation framework. Further analysis of these connectivity patterns, described below, confirmed this hypothesis.

**Figure 8:**
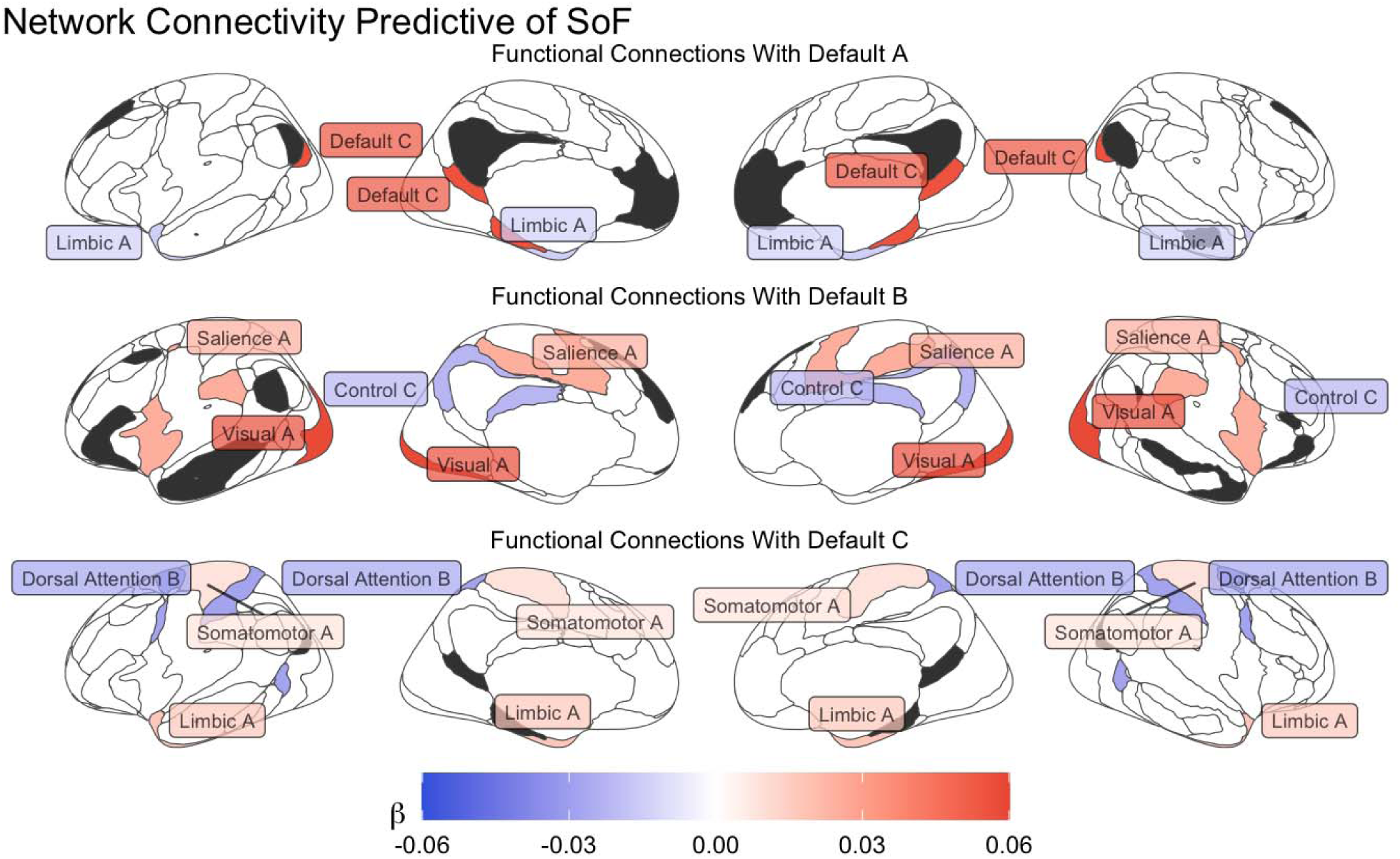
Functional network connectivity predictive of individual speeds of forgetting. The connectivity values are organized according to the DMN subnetwork they refer to (in dark gray); The color of each network reflects the value of the corresponding β weight of its functional connectivity.

**Table 1:** List of network connections predictive of individual SoF values, together with their β weights, in decreasing order.

| Network Connectivity | Regressor Weight ($\beta$) |
| --- | --- |
| Default A to Default C | 0.055810 |
| Default A to Limbic A | -0.014422 |
| Default B to Control C | -0.022168 |
| Default B to Salience / Ventral Attention A | 0.025213 |
| Default B to Visual Central (Visual A) | 0.058361 |
| Default C to Dorsal Attention B | -0.027954 |
| Default C to Limbic A | 0.017076 |
| Default C to Somatomotor A | 0.008430 |

The predictions from this model were then compared to the observed data. Figure 9A plots the predicted speeds of forgetting against the observed values estimated from the adaptive paired-associate learning task. The final model’s predictions have a mean correlation of *r*(33) = 0.77 (*p* < 0.001) with the observed data.

**Figure 9:**
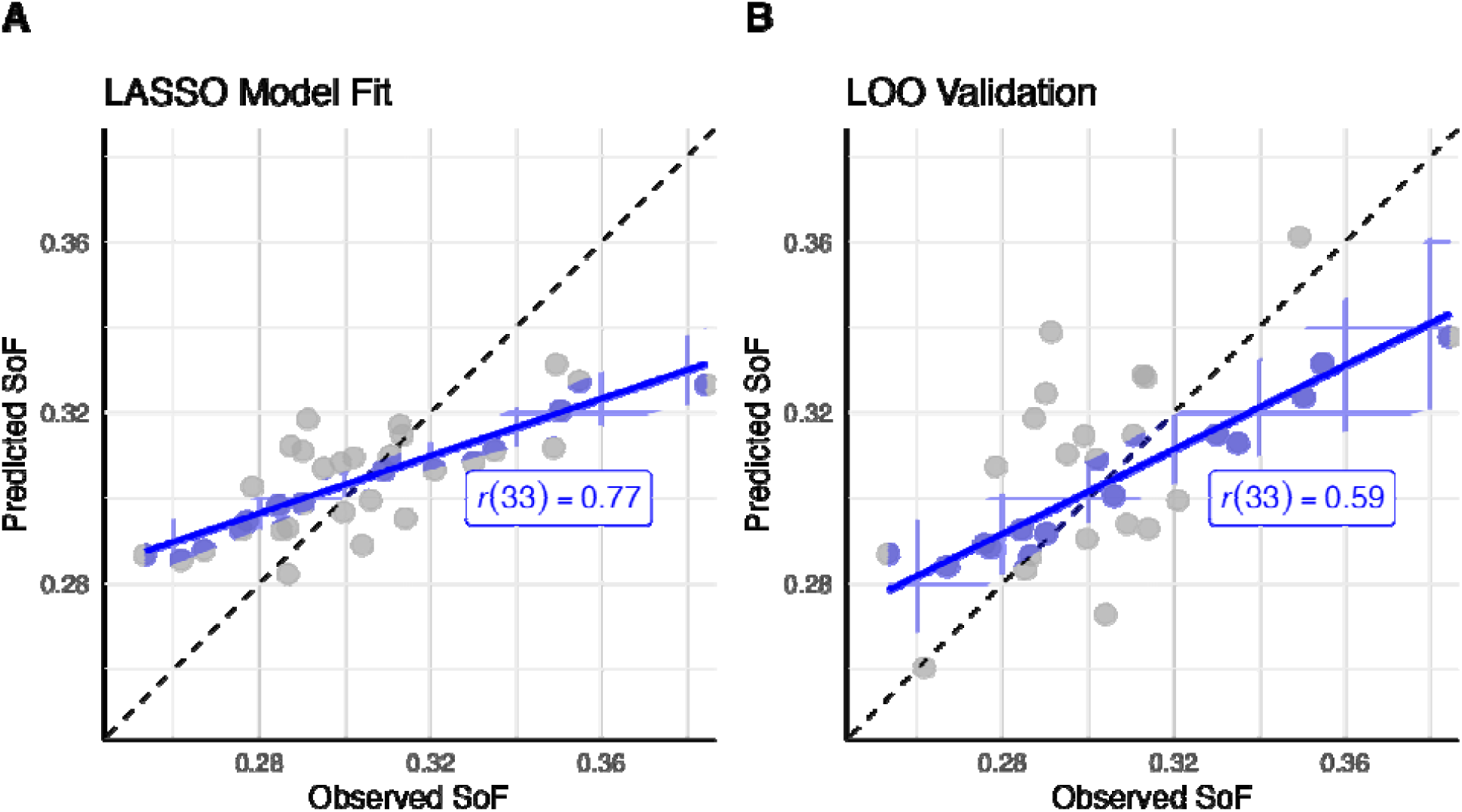
(A) Observed vs. predicted SoF values across all 33 participants, using the eight regressors identified by the LASSO Model. (B) Leave One Out (LOO) validation of the model, where each participant’s SoF value is predicted from beta weights fit to the remaining 32 individuals. In both plots, each dot represents a single participant; the black dashed line is the identity line, while the blue line and ribbon represent the linear regression estimate and its standard error.

### Validation of the LASSO Results

Given the relatively small sample size of this study, it is possible that the results of Figure 9A are partially due to overfitting and would not generalize. As a safeguard against this possibility, a Leave One Out cross-validation procedure was run on the final model. In this procedure, a prediction for each participant was generated by first fitting the beta weights of the eight regressors on the remaining 32 participants, and then by applying these weights to the specific connectivity values of the withheld participant. The results of this procedure are depicted in Figure 9B. As the panel shows, the model generalizes well to the held-out data, with the cross-validated predictions still having a significant correlation with the observed results [*r*(33) = 0.59 (*p* < 0.001)].

Because of its penalty term, LASSO might sometimes force the algorithm to select between collinear predictors at random, providing an inadequate and partial picture of which regressors are indeed essential [119]. If this were the case, other equally valid predictors could have been overlooked. A simple test to exclude this possibility is to remove the 8 connections from the original matrix of 45 regressors **X**, and re-run the LASSO procedure. If the procedure converges on a different set of features, then other important regressors were accidentally excluded. Alternatively, if no other regressors are left that can account for variability in the speed of forgetting, then it would be impossible for the cross-validation procedure to find a satisfying value of λ that allows for generalizable results. When applied to our data, this test shows that the cross-validation procedure does *not* converge anymore. Specifically, it selects a large hyperparameter value (λ = 0.0103) under whose constraints all regressors are set to zero; under these conditions, the resulting linear model reduces to the intercept and fails to capture any individual differences in forgetting.

To ascertain whether the eight regressors are *consistently* linked to the SoF values, we also performed eight follow-up LASSO analyses, each time removing only one of the eight regressors from the list of features. The removal of the connection between the Default C and Visual A sub-networks (which has the largest weight in Table 1) resulted in the model failing to converge. In each of the other cases, the model still converged to a solution and each of these solutions included all of the remaining seven regressors, suggesting that they are specifically and consistently associated with SoF values. In three of these seven cases, LASSO compensated by including 1–3 additional connections to the list of regressors. Only one such connection was common to these augmented solutions: the connection Default B and Visual B. Like one of our regressors, this connection links a DMN region with a visual one (thus suggesting a similar role) and also belongs to the Storage network group.

Taken together, these tests confirm that the eight regressor values identified by the LASSO model are reliably and consistently predictive of individual differences in SoF. However, since the number of predictors in the model (*p* = 8) remains proportionally large when compared to the number of observations (*n* = 33), it is possible that the surviving regressors are highly collinear, which would suggest that all connectivity values are similarly related to the speed of forgetting. To exclude this hypothesis, the eight regressors in the model were examined using the Variance Inflation Factor (VIF) metric as implemented in R’s *car* package [120]. The VIF of a regressor is the ratio of the variance of its estimate (that is, its β weight) in the full model over its variance in a reduced model that includes only that regressor. As a rule of thumb, VIF factors > 10 indicate collinearity [121]. This analysis revealed that all of the predictors had a VIF value < 2.5, which suggests that they contributed independently to an individual speed of forgetting.

Finally, a series of permutation tests was also conducted to examine whether any distribution of SoF values could be generated by an appropriate mixing of the full set of 45 regressors. To exclude this possibility, the LASSO procedure was repeated after randomly shuffling the vector of SoF values 1,000 times. In most cases (56.3%), the LASSO procedure failed to converge altogether. While possible linear solutions were identified in the remaining 43.6% of the simulations, models that were comparable to our results in terms of fit to the data were found in only 2.9% of the cases. Thus, it is statistically unlikely (*p* < 0.029) that the relationship between the distribution of SoF values and the distributions of connectivity values is is coincidental

### Relative Importance of DMN Networks

Having confirmed the internal validity of the model, we can now further investigate the nature of its predictors. The connections in Table 1 can be thought of as weighted edges connecting nodes in a network, with the weights corresponding to their β values; therefore, they can be analyzed using network-level measures. In particular, it is possible to compute the relative contribution of each of the Yeo networks as the *importance* of the corresponding node, which is the L1 norm of all of its connections to other nodes. In this case, each node is one of the Yeo networks, and its importance is the sum of the absolute value of the regressors that connect it to other Yeo networks in Table 1. As a first step, we computed the importance of each of the three DMN subnetworks (Default A, B, and C in Figure 3). The results are shown in Figure 10A.

**Figure 10:**
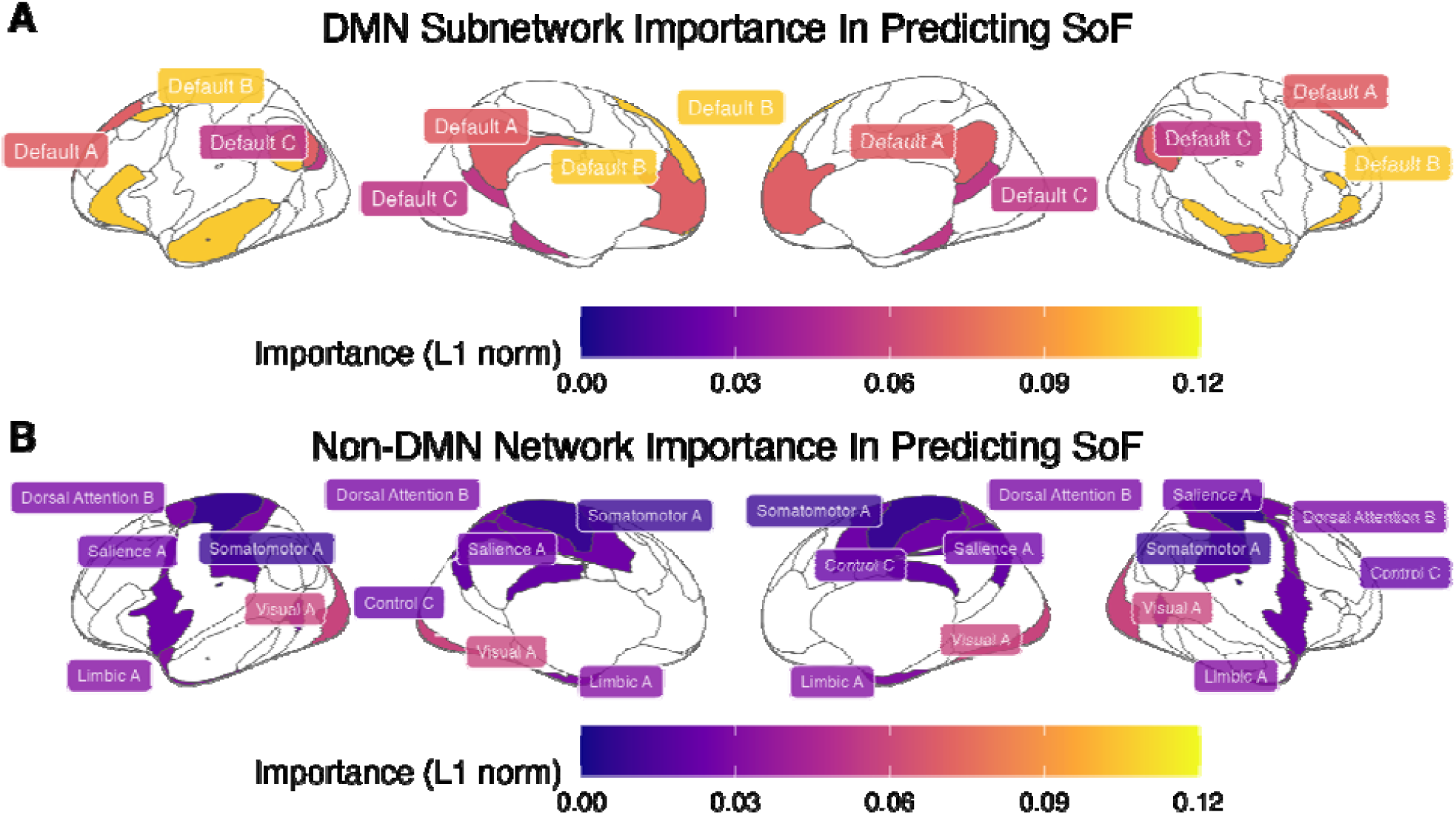
(A) Relative importance of each DMN subnetwork, calculated as the L1 norm of its connections. (B) Relative importance of each non-DMN network, calculated as the L1 norm of its connections.

### Relative Importance of Storage vs. Retrieval Networks

Figure 10B illustrates the importance of each of the six non-DMN subnetworks networks identified by the LASSO model (Somatomotor A, Visual A, Salience A, Control C, Limbic A, and Dorsal Attention B). As noted above, all of them belong to the storage-related group. Even accounting for the fact that our classification contains more storage networks (eight, accounting for 24 connections to the DMN subnetworks) than retrieval networks (six, for a total of 18 connections), the probability that *all* eight regressors would involve only storage networks is *p* < 0.003.

If the storage degradation hypothesis is correct, we would expect that the storage-related networks of Figure 4 would not only be selected more frequently, but would also have significantly greater importance than the retrieval-related networks. This is indeed confirmed by Figure 11, which illustrates the distribution of the importance scores of *all* of the non-DMN networks. A Wilcoxon test further confirmed that, even when all of the non-DMN networks were included (using a regressor value of zero for those eliminated by LASSO), the mean importance of the storage networks (*M*= 0.021, *SD* = 0.019) was significantly greater than zero (*t*(7) = 3.179, *p* = 0.016) and, as a consequence, greater than the mean across the retrieval networks (which is zero).

**Figure 11:**
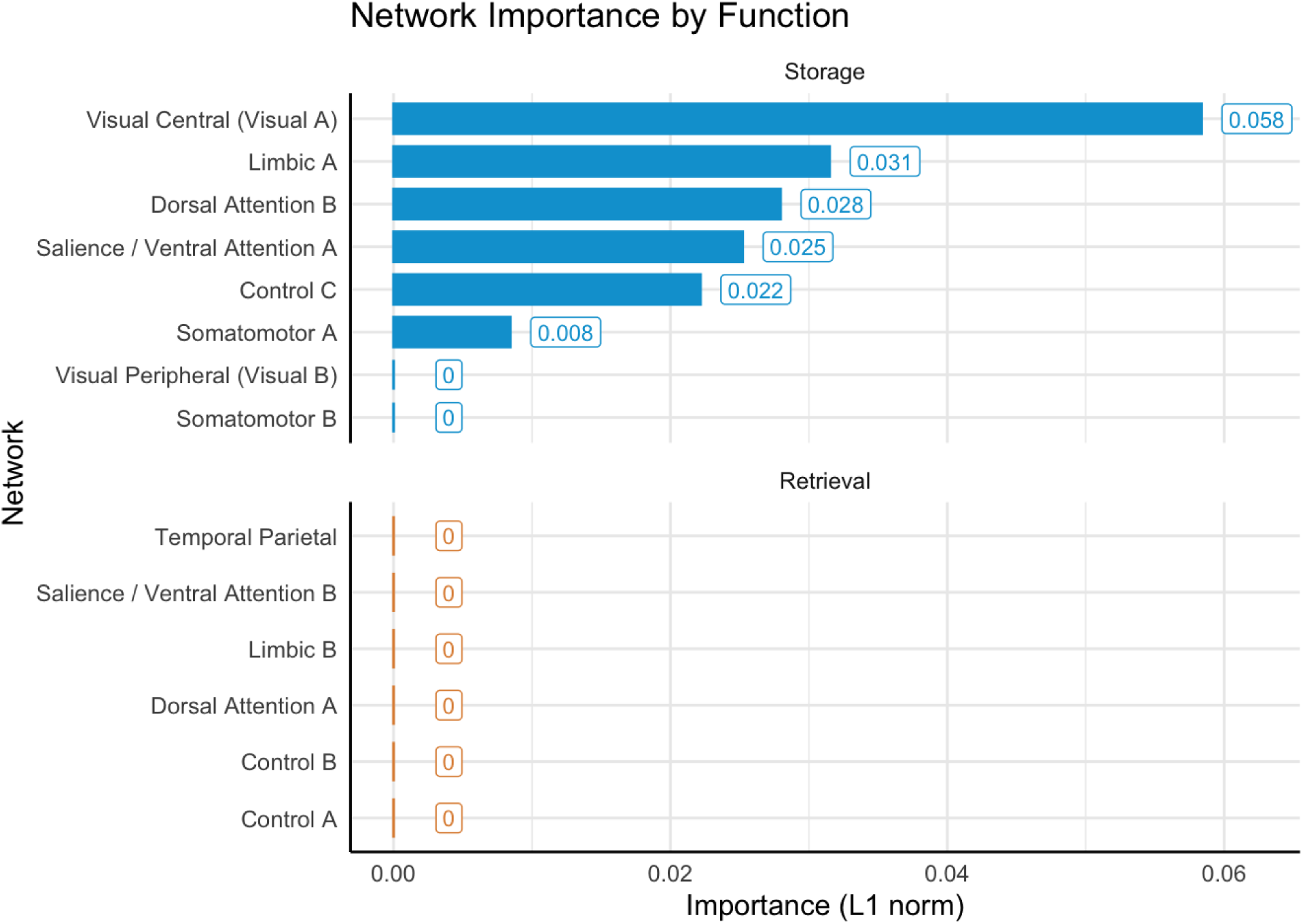
Connectivity importance of storage and retrieval networks to predict memory performance.

## Discussion

In this paper, we provided evidence that an individual’s speed of forgetting, operationalized as the decay intercept parameter [64] averaged over items, can be predicted using a distributed pattern of functional connectivity obtained from resting-state (or task-free) brain connectivity measures. This set of connections was identified through LASSO regression, demonstrating remarkable predictive accuracy while also reducing the full connectome to a manageable set of predictors. Functionally, these findings imply that forgetting primarily involves connectivity within the DMN and between the DMN and the storage-related networks, providing evidence in favor of the storage degradation hypothesis over the retrieval failure account.

### Limitations

While these findings are encouraging, it is important to recognize some of their inherent limitations. Although the use of a computational model allows a finer-grained understanding of the underlying memory process, our estimates of individual speeds of forgetting are dependent on the model’s default parameters for perceptual and motor processing. Although the values of these parameters are approximately correct for the current population (as shown by the model’s ability to fit the data, as illustrated in the Figure 6 and in the Supplementary Materials), different estimates might be necessary to correctly measure memory in different populations.

Most importantly, the data comes from a small number of fairly homogeneous participants who studied a relatively small set of foreign vocabulary items, which potentially limits external validity. Thus, future studies should expand the sample size and explore other sets of study materials. The limited sample size also prevents us from entirely ruling out a role for retrieval failure. It is possible, for example, that the contribution of retrieval networks is significant and consistent but not visible in our limited sample. It is also possible that the relative contributions of storage degradation and retrieval failure are mediated by external factors, such as age and clinical conditions. Thus, storage degradation may be the main contributor to forgetting in healthy young adults (our sample’s population), but retrieval failure might play a greater role in elderly adults or in patients affected by dementia. Once more, future studies will be needed to further examine these alternatives.

A related limitation concerns the interpretability of the specific connectivity results in Figure 8. It is impossible to judge how much the connectivity between two networks affects forgetting, because its final effects depend on both the size of its beta value and the value of the correlation. Even more strikingly, it is impossible to judge whether a higher or lower beta value for a functional connection translates to higher or lower forgetting, because the effect depends on the direction of the original functional connectivity. For example, if two regions were *anti*-correlated but their functional connectivity had a positive weight, an increase in functional connectivity would decrease, instead of increase, the predicted speed of forgetting. In a previous study by Yang et al. [122], the authors worked around this problem by multiplying the beta weight of a connection by the sign of its mean, group-level connectivity value, so that the sign of the product would correctly reflect the direction of the effect and the absolute value would predict the true predictive strength. If applied to our data, this approach yields the results illustrated in Figure 12.

**Figure 12:**
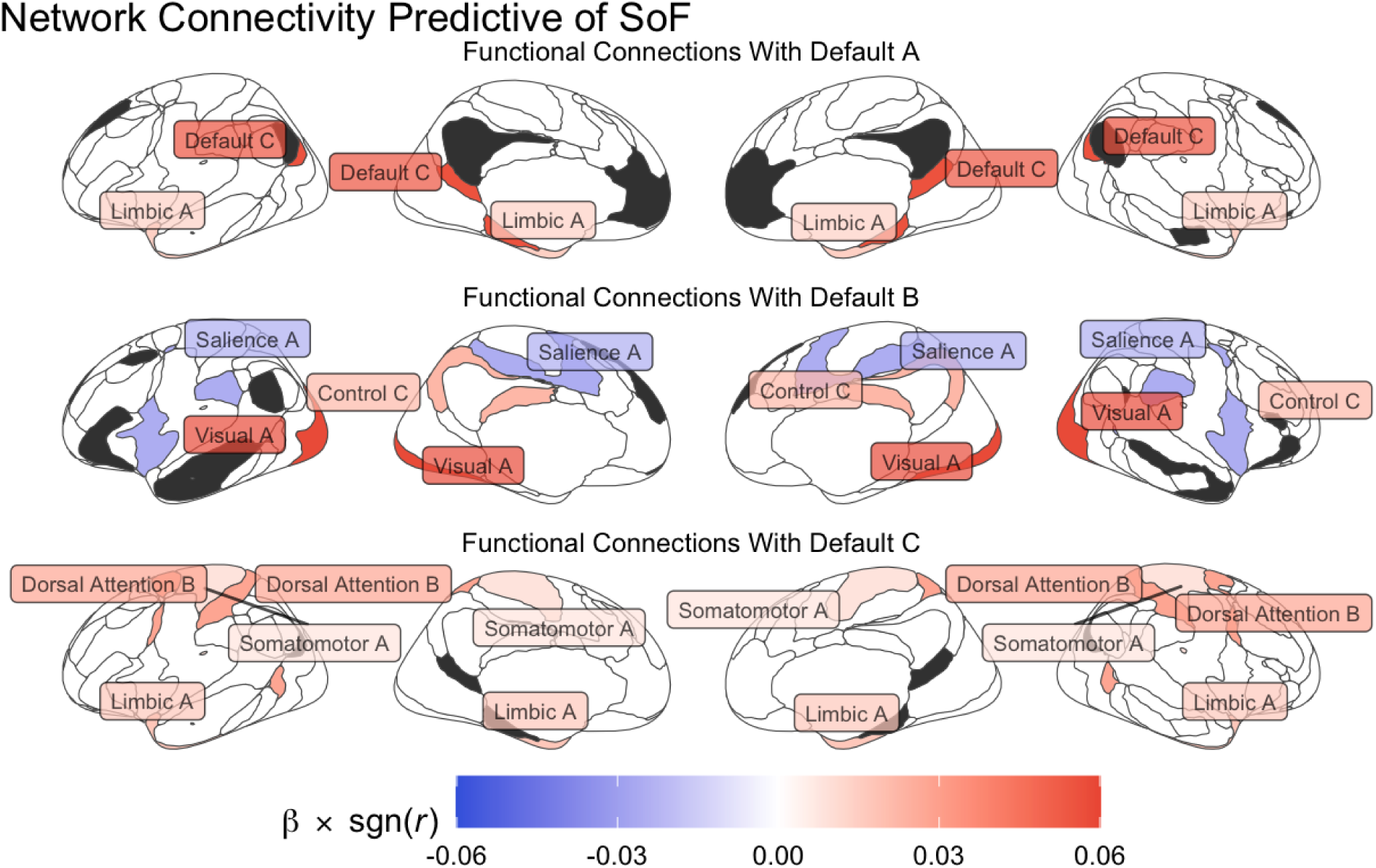
Interpretable functional connectivity predictive of individual speeds of forgetting, organized according to the DMN subnetwork they refer to (in dark gray). The color of each network reflects the value of the corresponding β weight multiplied by its mean functional connectivity; positive values are associated with increases (and negative values with decrease) in the Speed of Forgetting.

The results in Figure 12 suggest that *greater* connectivity at rest between storage-related networks and the DMN are associated with increased forgetting—with the only exception of the connectivity between the salience network and the DMN. This observation runs against the notion that spontaneous brain activity at rest supports consolidation; in turn, this suggests two possible interpretations. The first is that, while a tonic amount of functional activity does support memory consolidation and maintenance, too much of it might result in greater interference during memory reactivation and replay.

The second interpretation is that greater resting-state activity between these networks might be associated with shallower *encoding* of the initial engram, thus setting the stage for faster forgetting. For example, noisier spontaneous activity in the background might limit the interference with the recruitment of neurons to create the initial representation ensemble.

Indeed, it is possible that individual differences in the speed of forgetting might be ascribed due to individual differences in the depth of encoding. Although the computational model used in this research can successfully track differences in retention curves, it is agnostic about their underlying causes, leaving open the possibility that differences in forgetting are due to encoding. This is also consistent with a recent study [123] in which the authors experimentally manipulated the levels of processings [124,125] associated with different items and tracked their recall at different intervals and found that shallower processing at encoding indeed resulted in faster-declining retention curves.

Although both hypotheses are worth pursuing in future research, we should note that any interpretation of our results relies on having reliable estimates of the sign of group-level mean functional connectivity. Thus, given the limited size of our sample and the intrinsic limits of interpreting functional connectivity [126,127], the results in Figure 12 should be taken with caution.

### Considerations

These limitations notwithstanding, we believe it is important to stress four characteristics of our findings. First, they are based on a measure that directly quantifies forgetting, a latent and unobservable process, through computational phenotyping; this approach provides an advantage over simpler behavioral tests of memory performance. As noted in the introduction, forgetting is a *process* [3,7] and there is no consensus on how long after encoding a memory should be tested to precisely quantify retention. Since long-term memory retention spans from seconds to decades, it is difficult to test it in a laboratory setting. Consequently, previous studies have relied on individuals exhibiting dramatic differences in memory function, such as amnesic patients (such as patients suffering from medial temporal lobe injury [128–130]), while most laboratory experiments focus on relatively short retention intervals. Two participants, however, might have comparable accuracy-based performance a few minutes after studying but vastly different performances weeks later. In contrast, our findings rely on a novel metric, the speed of forgetting, that quantifies the rate at which memories are forgotten, and is better suited for capturing forgetting and retention at different intervals and, ultimately, for making inferences as to where forgetting might happen in the brain.

Second, it is worth noting that our findings could be applied in the opposite direction: to decode an individual’s speed of forgetting from distributed patterns of connectivity using a standard brain parcellation. Such an approach could be used, for example, to detect the early onset of abnormal forgetting in individuals with known memory pathologies and cognitive degenerative diseases. Most importantly, this finding suggests the possibility of decoding model parameters directly from resting state data, rather than from a combination of specific tasks. This procedure could play an important role in the development of high-fidelity idiographic models. This procedure also has the benefit of showing reflections of parameters in brain connectivity network regions, shedding light on a more holistic and comprehensive way of doing functional anatomy research on human cognition.

Third, our results are broadly consistent with the existing literature. As predicted, we did find that individual differences in forgetting covary with connectivity with and within the DMN. Not only have these sets of networks been theorized to play a role in memory replay and consolidation [10], but their connectivity with the salience network and the dorsal attention network (both of which were identified in our analysis) are altered in pathologies that negatively affect long-term memory function, such as Alzheimer’s Disease [12], mild traumatic brain injury [131] and depression [132]. Our findings are also consistent with previous studies that have experimentally tested the root causes of drug-induced amnesia [133,134], which similarly strongly favor the storage degradation hypothesis.

Finally, our results are also consistent with the findings of a previous study that previously attempted to identify the neural underpinnings of forgetting using the *same* model-based approach but with a different neuroimaging modality—EEG instead of fMRI [80]. In that study, a significant positive correlation between spectral power in the beta band (12-20 Hz) and the speed of forgetting was observed distributing at middle frontal (AF3 and AF4), bilateral posterior (channel O1, O2), and parietal (channel P8) regions; these spatial locations are consistent with the cortical regions of the DMN subnetworks A and B (see Figures 10 and 11).

## Conclusions

In summary, although further research will be needed to clarify the specific mechanisms of forgetting, our findings do provide a preliminary answer to the question of *where* memories are lost—that is, in the connections between the DMN and the sensory networks that are involved in the encoding, re-experience, and consolidation of a memory. Thus, our results imply that forgetting can be interpreted as a storage degradation process, rather than a retrieval access problem, and that future clinical and neuroscientific research should focus on the storage networks as a target to better understand, improve, or support memory function.

## Acknowledgments

This work was supported by grant N00014-17-1-2607 from the Office of Naval Research to CP and by grant FA9550-19-1-0299 from the Air Force Office of Scientific Research to AS.

## Footnotes

1 Unfortunately, the trial-level data from two of our participants have been lost, and could not be entered into this analysis.

## References

1. Eichenbaum H. Declarative memory: insights from cognitive neurobiology. Annu Rev Psychol. 1997;48: 547–572.

2. Feld GB, Born J. Sculpting memory during sleep: concurrent consolidation and forgetting. Current Opinion in Neurobiology. 2017;44: 20–27.

3. Hardt O, Nader K, Nadel L. Decay happens: the role of active forgetting in memory. Trends Cogn Sci. 2013;17: 111–120.

4. Guskjolen A, Cembrowski MS. Engram neurons: Encoding, consolidation, retrieval, and forgetting of memory. Mol Psychiatry. 2023;28: 3207–3219.

5. Shiffrin RM. Forgetting: Trace Erosion or Retrieval Failure? Science. 1970;168: 1601– 1603.

6. Spear NE. Forgetting as retrieval failure. Animal memory. 1971; 45–109.

7. Davis RL, Zhong Y. The biology of forgetting—A perspective. Neuron. 2017;95: 490–503.

8. Ryan TJ, Frankland PW. Forgetting as a form of adaptive engram cell plasticity. Nat Rev Neurosci. 2022;23: 173–186.

9. Frankland PW, Köhler S, Josselyn SA. Hippocampal neurogenesis and forgetting. Trends Neurosci. 2013;36: 497–503.

10. Pezzulo G, Zorzi M, Corbetta M. The secret life of predictive brains: what’s spontaneous activity for? Trends Cogn Sci. 2021;25: 730–743.

11. Jin M, Pelak VS, Cordes D. Aberrant default mode network in subjects with amnestic mild cognitive impairment using resting-state functional MRI. Magn Reson Imaging. 2012;30: 48–61.

12. Qi H, Liu H, Hu H, He H, Zhao X. Primary Disruption of the Memory-Related Subsystems of the Default Mode Network in Alzheimer’s Disease: Resting-State Functional Connectivity MRI Study. Front Aging Neurosci. 2018;10: 344.

13. Alfei JM, Miller RR, Ryan TJ, Urcelay GP. Rethinking memory impairments: Retrieval failure. Psychol Rev. 2025. doi:10.1037/rev0000538

14. Penfield W. Engrams in the human brain. Mechanisms of memory. Proc R Soc Med. 1968;61: 831–840.

15. Curot J, Busigny T, Valton L, Denuelle M, Vignal J-P, Maillard L, et al. Memory scrutinized through electrical brain stimulation: A review of 80 years of experiential phenomena. Neurosci Biobehav Rev. 2017;78: 161–177.

16. Sjöberg RL. Brain stimulation and elicited memories. Acta Neurochir (Wien). 2023;165: 2737–2745.

17. Loftus EF, Loftus GR. On the permanence of stored information in the human brain. Am Psychol. 1980;35: 409–420.

18. Roy DS, Arons A, Mitchell TI, Pignatelli M, Ryan TJ, Tonegawa S. Memory retrieval by activating engram cells in mouse models of early Alzheimer’s disease. Nature. 2016;531: 508–512.

19. Swanson CJ, Zhang Y, Dhadda S, Wang J, Kaplow J, Lai RYK, et al. A randomized, double-blind, phase 2b proof-of-concept clinical trial in early Alzheimer’s disease with lecanemab, an anti-Aβ protofibril antibody. Alzheimers Res Ther. 2021;13: 80.

20. Cummings J, Apostolova L, Rabinovici GD, Atri A, Aisen P, Greenberg S, et al. Lecanemab: Appropriate Use Recommendations. J Prev Alzheimers Dis. 2023;10: 362–377.

21. Ranganath C, Heller A, Cohen MX, Brozinsky CJ, Rissman J. Functional connectivity with the hippocampus during successful memory formation. Hippocampus. 2005;15: 997–1005.

22. Fernández G, Effern A, Grunwald T, Pezer N, Lehnertz K, Dümpelmann M, et al. Real-time tracking of memory formation in the human rhinal cortex and hippocampus. Science. 1999;285: 1582–1585.

23. Poo M-M, Pignatelli M, Ryan TJ, Tonegawa S, Bonhoeffer T, Martin KC, et al. What is memory? The present state of the engram. BMC Biol. 2016;14: 40.

24. Josselyn SA, Tonegawa S. Memory engrams: Recalling the past and imagining the future. Science. 2020;367: eaaw4325.

25. Danker JF, Anderson JR. The ghosts of brain states past: remembering reactivates the brain regions engaged during encoding. Psychol Bull. 2010;136: 87–102.

26. Huang Q, Xiao Z, Yu Q, Luo Y, Xu J, Qu Y, et al. Replay-triggered brain-wide activation in humans. Nat Commun. 2024;15: 7185.

27. Ranganath C, Ritchey M. Two cortical systems for memory-guided behaviour. Nat Rev Neurosci. 2012;13: 713–726.

28. Kaefer K, Stella F, McNaughton BL, Battaglia FP. Replay, the default mode network and the cascaded memory systems model. Nat Rev Neurosci. 2022;23: 628–640.

29. Raichle ME, Snyder AZ. A default mode of brain function: a brief history of an evolving idea. Neuroimage. 2007;37: 1083–90; discussion 1097-9.

30. Sibert C, Hake HS, Stocco A. The Structured Mind at Rest: Low-Frequency Oscillations Reflect Interactive Dynamics Between Spontaneous Brain Activity and a Common Architecture for Task Control. Front Neurosci. 2022;16: 832503.

31. Kim H. Dissociating the roles of the default-mode, dorsal, and ventral networks in episodic memory retrieval. Neuroimage. 2010;50: 1648–1657.

32. Philippi CL, Tranel D, Duff M, Rudrauf D. Damage to the default mode network disrupts autobiographical memory retrieval. Soc Cogn Affect Neurosci. 2015;10: 318–326.

33. Menon V. 20 years of the default mode network: A review and synthesis. Neuron. 2023;111: 2469–2487.

34. Petrides M. The mid-ventrolateral prefrontal cortex and active mnemonic retrieval. Neurobiol Learn Mem. 2002;78: 528–538.

35. Badre D, Wagner AD. Left ventrolateral prefrontal cortex and the cognitive control of memory. Neuropsychologia. 2007;45: 2883–2901.

36. Thompson-Schill SL, D’Esposito M, Aguirre GK, Farah MJ. Role of left inferior prefrontal cortex in retrieval of semantic knowledge: a reevaluation. Proc Natl Acad Sci U S A. 1997;94: 14792–14797.

37. Danker JF, Gunn P, Anderson JR. A rational account of memory predicts left prefrontal activation during controlled retrieval. Cereb Cortex. 2008;18: 2674–2685.

38. Borst JP, Anderson JR. Using model-based functional MRI to locate working memory updates and declarative memory retrievals in the fronto-parietal network. Proc Natl Acad Sci U S A. 2013;110: 1628–1633.

39. Danker JF, Fincham JM, Anderson JR. The neural correlates of competition during memory retrieval are modulated by attention to the cues. Neuropsychologia. 2011;49: 2427–2438.

40. Lepage M, Ghaffar O, Nyberg L, Tulving E. Prefrontal cortex and episodic memory retrieval mode. Proc Natl Acad Sci U S A. 2000;97: 506–511.

41. Chua EF, Schacter DL, Sperling RA. Neural correlates of metamemory: a comparison of feeling-of-knowing and retrospective confidence judgments. J Cogn Neurosci. 2009;21: 1751–1765.

42. Modirrousta M, Fellows LK. Medial prefrontal cortex plays a critical and selective role in “feeling of knowing” meta-memory judgments. Neuropsychologia. 2008;46: 2958–2965.

43. Schnyer DM, Verfaellie M, Alexander MP, LaFleche G, Nicholls L, Kaszniak AW. A role for right medial prefrontal cortex in accurate feeling-of-knowing judgments: evidence from patients with lesions to frontal cortex. Neuropsychologia. 2004;42: 957–966.

44. Turner MS, Cipolotti L, Yousry TA, Shallice T. Confabulation: damage to a specific inferior medial prefrontal system. Cortex. 2008;44: 637–648.

45. Benson DF, Djenderedjian A, Miller BL, Pachana NA, Chang L, Itti L, et al. Neural basis of confabulation. Neurology. 1996;46: 1239–1243.

46. Raichle ME. The brain’s default mode network. Annu Rev Neurosci. 2015;38: 433–447.

47. McClelland JL, McNaughton BL, O’Reilly RC. Why there are complementary learning systems in the hippocampus and neocortex: insights from the successes and failures of connectionist models of learning and memory. Psychol Rev. 1995;102: 419.

48. O’Reilly RC, Bhattacharyya R, Howard MD, Ketz N. Complementary learning systems. Cogn Sci. 2014;38: 1229–1248.

49. Alvarez P, Squire LR. Memory consolidation and the medial temporal lobe: a simple network model. Proc Natl Acad Sci U S A. 1994;91: 7041–7045.

50. Zhang M, Nathaniel U, Savill N, Smallwood J, Jefferies E. Intrinsic connectivity of left ventrolateral prefrontal cortex predicts individual differences in controlled semantic retrieval. Neuroimage. 2022;246: 118760.

51. Loftus GR. On interpretation of interactions. Mem Cognit. 1978;6: 312–319.

52. Brady TF, Robinson MM, Williams JR, Wixted JT. Measuring memory is harder than you think: How to avoid problematic measurement practices in memory research. Psychon Bull Rev. 2023;30: 421–449.

53. Patzelt EH, Hartley CA, Gershman SJ. Computational phenotyping: Using models to understand individual differences in personality, development, and mental illness. Personal Neurosci. 2018;1: e18.

54. Schurr R, Reznik D, Hillman H, Bhui R, Gershman SJ. Dynamic computational phenotyping of human cognition. Nat Hum Behav. 2024;8: 917–931.

55. Sense F, Behrens F, Meijer RR, van Rijn H. An individual’s rate of forgetting is stable over time but differs across materials. Top Cogn Sci. 2016;8: 305–321.

56. Xu Y, Stocco A. Recovering Reliable Idiographic Biological Parameters from Noisy Behavioral Data: the Case of Basal Ganglia Indices in the Probabilistic Selection Task. Comput Brain Behav. 2021; 1–17.

57. Marek S, Tervo-Clemmens B, Calabro FJ, Montez DF, Kay BP, Hatoum AS, et al. Reproducible brain-wide association studies require thousands of individuals. Nature. 2022;603: 654–660.

58. Daw ND. Trial-by-trial data analysis using computational models. Decision making, affect, and learning: Attention and performance XXIII. 2011;23: 3–38.

59. Collins AGE. The Tortoise and the Hare: Interactions between Reinforcement Learning and Working Memory. J Cogn Neurosci. 2018;30: 1422–1432.

60. White CN, Congdon E, Mumford JA, Karlsgodt KH, Sabb FW, Freimer NB, et al. Decomposing decision components in the stop-signal task: a model-based approach to individual differences in inhibitory control. J Cogn Neurosci. 2014;26: 1601–1614.

61. White CN, Curl RA, Sloane JF. Using decision models to enhance investigations of individual differences in cognitive neuroscience. Front Psychol. 2016;7: 81.

62. Wagenmakers E-J, Krypotos A-M, Criss AH, Iverson G. On the interpretation of removable interactions: a survey of the field 33 years after Loftus. Mem Cognit. 2012;40: 145–160.

63. Anderson JR, Schooler LJ. Reflections of the Environment in Memory. Psychol Sci. 1991;2: 396–408.

64. Pavlik PI Jr, Anderson JR. Practice and forgetting effects on vocabulary memory: An activation-based model of the spacing effect. Cogn Sci. 2005;29: 559–586.

65. Pavlik PI, Anderson JR. Using a model to compute the optimal schedule of practice. J Exp Psychol Appl. 2008;14: 101–117.

66. Sense F, Meijer RR, van Rijn H. Exploration of the Rate of Forgetting as a Domain-Specific Individual Differences Measure. Frontiers in Education. 2018;3: 112.

67. Anderson JR. How Can the Human Mind Occur in the Physical Universe? Oxford University Press; 2007.

68. Anderson JR, Bothell D, Byrne MD, Douglass S, Lebiere C, Qin Y. An integrated theory of the mind. Psychol Rev. 2004;111: 1036–1060.

69. Stocco A, Rice P, Thomson R, Smith B, Morrison D, Lebiere C. An Integrated Computational Framework for the Neurobiology of Memory Based on the ACT-R Declarative Memory System. Computational Brain & Behavior. 2023. doi:10.1007/s42113-023-00189-y

70. van Rijn D, Maanen L, Woudenberg MV. Passing the test: Improving learning gains by balancing spacing and testing effects. Proceedings of the 9th. 2009; 108–114.

71. van der Velde M, Sense F, Borst JP, van Rijn H. Large-scale evaluation of cold-start mitigation in adaptive fact learning: Knowing “what” matters more than knowing “who.” User Model User-adapt Interact. 2024. doi:10.1007/s11257-024-09401-5

72. Wilschut T, Sense F, van Rijn H. Speaking to remember: Model-based adaptive vocabulary learning using automatic speech recognition. Comput Speech Lang. 2024;84: 101578.

73. Nadel L, Samsonovich A, Ryan L, Moscovitch M. Multiple trace theory of human memory: computational, neuroimaging, and neuropsychological results. Hippocampus. 2000;10: 352–368.

74. Moscovitch M, Rosenbaum RS, Gilboa A, Addis DR, Westmacott R, Grady C, et al. Functional neuroanatomy of remote episodic, semantic and spatial memory: a unified account based on multiple trace theory. J Anat. 2005;207: 35–66.

75. Rubin DC, Wenzel AE. One hundred years of forgetting: A quantitative description of retention. Psychol Rev. 1996;103: 734–760.

76. van der Velde M, Sense F, Borst JP, van Rijn H. Alleviating the Cold Start Problem in Adaptive Learning using Data-Driven Difficulty Estimates. Computational Brain & Behavior. 2021;4: 231–249.

77. Sense F, Maaß S, Gluck K, van Rijn H. Within-Subject Performance on a Real-Life, Complex Task and Traditional Lab Experiments: Measures of Word Learning, Raven Matrices, Tapping, and CPR. J Cogn. 2019;2: 12.

78. Sense F, Behrens F, Meijer RR, van Rijn H. Stability of Individual Parameters in a Model of Optimal Fact Learning. Proceedings of the 13th International Conference on Cognitive Modeling. 2015. pp. 136–141.

79. Shrout PE, Lane SP. Psychometrics. In: Mehl MR, editor. Handbook of research methods for studying daily life , (pp. New York, NY, US: The Guilford Press, xxvii; 2012. pp. 302– 320.

80. Zhou P, Sense F, van Rijn H, Stocco A. Reflections of idiographic long-term memory characteristics in resting-state neuroimaging data. Cognition. 2021;212: 104660.

81. James G, Witten D, Hastie T, Tibshirani R. Statistical Learning. In: James G, Witten D, Hastie T, Tibshirani R, editors. An Introduction to Statistical Learning: with Applications in R. New York, NY: Springer New York; 2013. pp. 15–57.

82. Braun U, Plichta MM, Esslinger C, Sauer C, Haddad L, Grimm O, et al. Test-retest reliability of resting-state connectivity network characteristics using fMRI and graph theoretical measures. Neuroimage. 2012;59: 1404–1412.

83. Stocco A, Sibert C, Steine-Hanson Z, Koh N, Laird JE, Lebiere CJ, et al. Analysis of the human connectome data supports the notion of a “Common Model of Cognition” for human and human-like intelligence across domains. Neuroimage. 2021;235: 118035.

84. Finn ES, Shen X, Scheinost D, Rosenberg MD, Huang J, Chun MM, et al. Functional connectome fingerprinting: identifying individuals using patterns of brain connectivity. Nat Neurosci. 2015;18: 1664–1671.

85. Waller L, Walter H, Kruschwitz JD, Reuter L, Müller S, Erk S, et al. Evaluating the replicability, specificity, and generalizability of connectome fingerprints. Neuroimage. 2017;158: 371–377.

86. Avery EW, Yoo K, Rosenberg MD, Greene AS, Gao S, Na DL, et al. Distributed Patterns of Functional Connectivity Predict Working Memory Performance in Novel Healthy and Memory-impaired Individuals. J Cogn Neurosci. 2020;32: 241–255.

87. Reineberg AE, Andrews-Hanna JR, Depue BE, Friedman NP, Banich MT. Resting-state networks predict individual differences in common and specific aspects of executive function. Neuroimage. 2015;104: 69–78.

88. Reineberg AE, Banich MT. Functional connectivity at rest is sensitive to individual differences in executive function: A network analysis. Hum Brain Mapp. 2016;37: 2959– 2975.

89. McGregor HR, Gribble PL. Functional connectivity between somatosensory and motor brain areas predicts individual differences in motor learning by observing. J Neurophysiol. 2017;118: 1235–1243.

90. Baldassarre A, Lewis CM, Committeri G, Snyder AZ, Romani GL, Corbetta M. Individual variability in functional connectivity predicts performance of a perceptual task. Proc Natl Acad Sci U S A. 2012;109: 3516–3521.

91. Persson J, Stening E, Nordin K, Söderlund H. Predicting episodic and spatial memory performance from hippocampal resting-state functional connectivity: Evidence for an anterior-posterior division of function. Hippocampus. 2018;28: 53–66.

92. Durrant S, Lewis PA. Memory consolidation: tracking transfer with functional connectivity. Curr Biol. 2009;19: R860–2.

93. Van den Broek GSE, Segers E, Van Rijn H, Takashima A, Verhoeven L. Effects of elaborate feedback during practice tests: Costs and benefits of retrieval prompts. J Exp Psychol Appl. 2019;25: 588–601.

94. Nelson TO, Dunlosky J. Norms of paired-associate recall during multitrial learning of Swahili-English translation equivalents. Memory. 1994;2: 325–335.

95 . Roediger HL 3rd, Butler AC. The critical role of retrieval practice in long-term retention. Trends Cogn Sci. 2011;15: 20–27.

96. Anderson JR. Retrieval of propositional information from long-term memory. Cogn Psychol. 1974;6: 451–474.

97. Anderson JR. How Can the Mind Occur in the Physical Universe? Oxford University Press; 2007.

98. Cox RW. AFNI: software for analysis and visualization of functional magnetic resonance neuroimages. Comput Biomed Res. 1996;29: 162–173.

99. Yeo BTT, Krienen FM, Sepulcre J, Sabuncu MR, Lashkari D, Hollinshead M, et al. The organization of the human cerebral cortex estimated by intrinsic functional connectivity. J Neurophysiol. 2011;106: 1125–1165.

100. Miyashita Y, Kameyama M, Hasegawa I, Fukushima T. Consolidation of visual associative long-term memory in the temporal cortex of primates. Neurobiol Learn Mem. 1998;70: 197–211.

101. Brawn TP, Fenn KM, Nusbaum HC, Margoliash D. Consolidation of sensorimotor learning during sleep. Learn Mem. 2008;15: 815–819.

102. Cuppone AV, Semprini M, Konczak J. Consolidation of human somatosensory memory during motor learning. Behav Brain Res. 2018;347: 184–192.

103. Krakauer JW, Shadmehr R. Consolidation of motor memory. Trends Neurosci. 2006;29: 58–64.

104. Squire LR, Genzel L, Wixted J, Morris RG. Memory consolidation. Cold Spring Harb Perspect Biol. 2015;7: a021766.

105. Casanova JP, Madrid C, Contreras M, Rodríguez M, Vasquez M, Torrealba F. A role for the interoceptive insular cortex in the consolidation of learned fear. Behav Brain Res. 2016;296: 70–77.

106. Bermudez-Rattoni F. The forgotten insular cortex: its role on recognition memory formation. Neurobiol Learn Mem. 2014;109: 207–216.

107. Andreano JM, Touroutoglou A, Dickerson BC, Barrett LF. Resting connectivity between salience nodes predicts recognition memory. Soc Cogn Affect Neurosci. 2017;12: 948–955.

108. Lu Y-T, Chang W-N, Chang C-C, Lu C-H, Chen N-C, Huang C-W, et al. Insula volume and salience network are associated with memory decline in Parkinson disease: Complementary analyses of voxel-based morphometry versus volume of interest. Parkinsons Dis. 2016;2016: 2939528.

109. Krauel K, Duzel E, Hinrichs H, Santel S, Rellum T, Baving L. Impact of emotional salience on episodic memory in attention-deficit/hyperactivity disorder: a functional magnetic resonance imaging study. Biol Psychiatry. 2007;61: 1370–1379.

110. Sporns O. The human connectome: a complex network. Ann N Y Acad Sci. 2011;1224: 109–125.

111. Cole MW, Ito T, Bassett DS, Schultz DH. Activity flow over resting-state networks shapes cognitive task activations. Nat Neurosci. 2016;19: 1718–1726.

112. Shen X, Finn ES, Scheinost D, Rosenberg MD, Chun MM, Papademetris X, et al. Using connectome-based predictive modeling to predict individual behavior from brain connectivity. Nat Protoc. 2017;12: 506–518.

113. Tibshirani R. Regression shrinkage and selection via the lasso. J R Stat Soc Series B Stat Methodol. 1996. Available: https://academic.oup.com/jrsssb/article-abstract/58/1/267/7027929

114. James G, Witten D, Hastie T, Tibshirani R. An Introduction to Statistical Learning. Springer US; 2013.

115. James G, Witten D, Hastie T, Tibshirani R, Taylor J. An Introduction to Statistical Learning: with Applications in Python. Springer Nature; 2023.

116. Myers CE, Interian A, Moustafa AA. A practical introduction to using the drift diffusion model of decision-making in cognitive psychology, neuroscience, and health sciences. Front Psychol. 2022;13: 1039172.

117. Fox MD, Snyder AZ, Vincent JL, Corbetta M, Van Essen DC, Raichle ME. The human brain is intrinsically organized into dynamic, anticorrelated functional networks. Proc Natl Acad Sci U S A. 2005;102: 9673–9678.

118. Jia J, Yu B. On model selection consistency of the Elastic Net when p Ill n. Stat Sin. 2010;20: 595–611.

119. Muthukrishnan R, Rohini R. LASSO: A feature selection technique in predictive modeling for machine learning. 2016 IEEE International Conference on Advances in Computer Applications (ICACA). IEEE; 2016. pp. 18–20.

120. Fox J, Weisberg S. An R Companion to Applied Regression. SAGE Publications; 2018.

121. Fox J, Monette G. Generalized Collinearity Diagnostics. J Am Stat Assoc. 1992;87: 178– 183.

122. Yang YC, Sibert C, Stocco A. Reliance on Episodic vs. Procedural Systems in Decision-Making Depends on Individual Differences in Their Relative Neural Efficiency. Computational Brain & Behavior. 2024. doi:10.1007/s42113-024-00203-x

123. Peng N, Logie RH, Della Sala S. Effect of levels-of-processing on rates of forgetting. Mem Cognit. 2025;53: 692–709.

124. Craik FIM. Levels of processing: past, present. and future? Memory. 2002;10: 305–318.

125. Lockhart RS, Craik FIM. Levels of processing: A retrospective commentary on a framework for memory research. Can J Psychol. 1990;44: 87–112.

126. Bastos AM, Schoffelen J-M. A tutorial review of functional connectivity analysis methods and their interpretational pitfalls. Front Syst Neurosci. 2015;9: 175.

127. Buckner R, Krienen FM, Yeo T. Opportunities and limitations of intrinsic functional connectivity MRI. Nat Neurosci. 2013;16: 832–837.

128. Scoville WB, Milner B. Loss of recent memory after bilateral hippocampal lesions. J Neurol Neurosurg Psychiatry. 1957;20: 11–21.

129. Corkin S. What’s new with the amnesic patient H.M.? Nat Rev Neurosci. 2002;3: 153– 160.

130. Gabrieli JD, Cohen NJ, Corkin S. The impaired learning of semantic knowledge following bilateral medial temporal-lobe resection. Brain Cogn. 1988;7: 157–177.

131. Andrews MJ, Salat DH, Milberg WP, McGlinchey RE, Fortier CB. Poor sleep and decreased cortical thickness in veterans with mild traumatic brain injury and post-traumatic stress disorder. Mil Med Res. 2024;11: 51.

132. Mulders PC, van Eijndhoven PF, Schene AH, Beckmann CF, Tendolkar I. Resting-state functional connectivity in major depressive disorder: A review. Neurosci Biobehav Rev. 2015;56: 330–344.

133. Hardt O, Wang S-H, Nader K. Storage or retrieval deficit: the yin and yang of amnesia. Learn Mem. 2009;16: 224–230.

134. Wang S-H, Finnie PSB, Hardt O, Nader K. Dorsal hippocampus is necessary for novel learning but sufficient for subsequent similar learning. Hippocampus. 2012;22: 2157–2170.

