## Supplementary Information for "Default Mode Network Connectivity Predicts Individual Differences in Long-Term Forgetting: Evidence for Storage Degradation, not Retrieval Failure"

### Goodness of Fit

To provide precise measures of the model’s fit to the data, we proceeded as follows. First, we collected individual φ values (Eq. 2) for each item studied by each participant. In the adaptive software used in this study, the final estimate of each item’s φ value is the value recorded for the last presentation of each stimulus. The average φ value across all items corresponds to the individual SOF values used in our functional connectivity analysis and reported in Figure 5.

The final φ value for each studied item was then used to compute the activation of its corresponding memory after the initial study (Figure 2). The activation of a memory at the time each test was computed using Eq. 1 and 2 and each cue’s presentation time. The corresponding predicted retrieval accuracy and response time for each trial were then computed from the memory’s activation using Eq. 3 and Eq. 4, respectively.

#### Predicted vs. Observed Accuracies

To compare predicted and observed accuracies, we first converted the observed participant responses to a binary coded dummy variable (0 = incorrect, 1 = correct). We then discretized the predicted retrieval probabilities into corresponding binary values, coded as 0 (if the probability of retrieval is ≤ 0.5) and 1 (if the probability > 0.5). Across all participants, the model’s accuracy in predicting the correct response was 0.71 (*p* < 0.001). Figure S1 illustrates the confusion matrices of the individual predicted and observed accuracies:

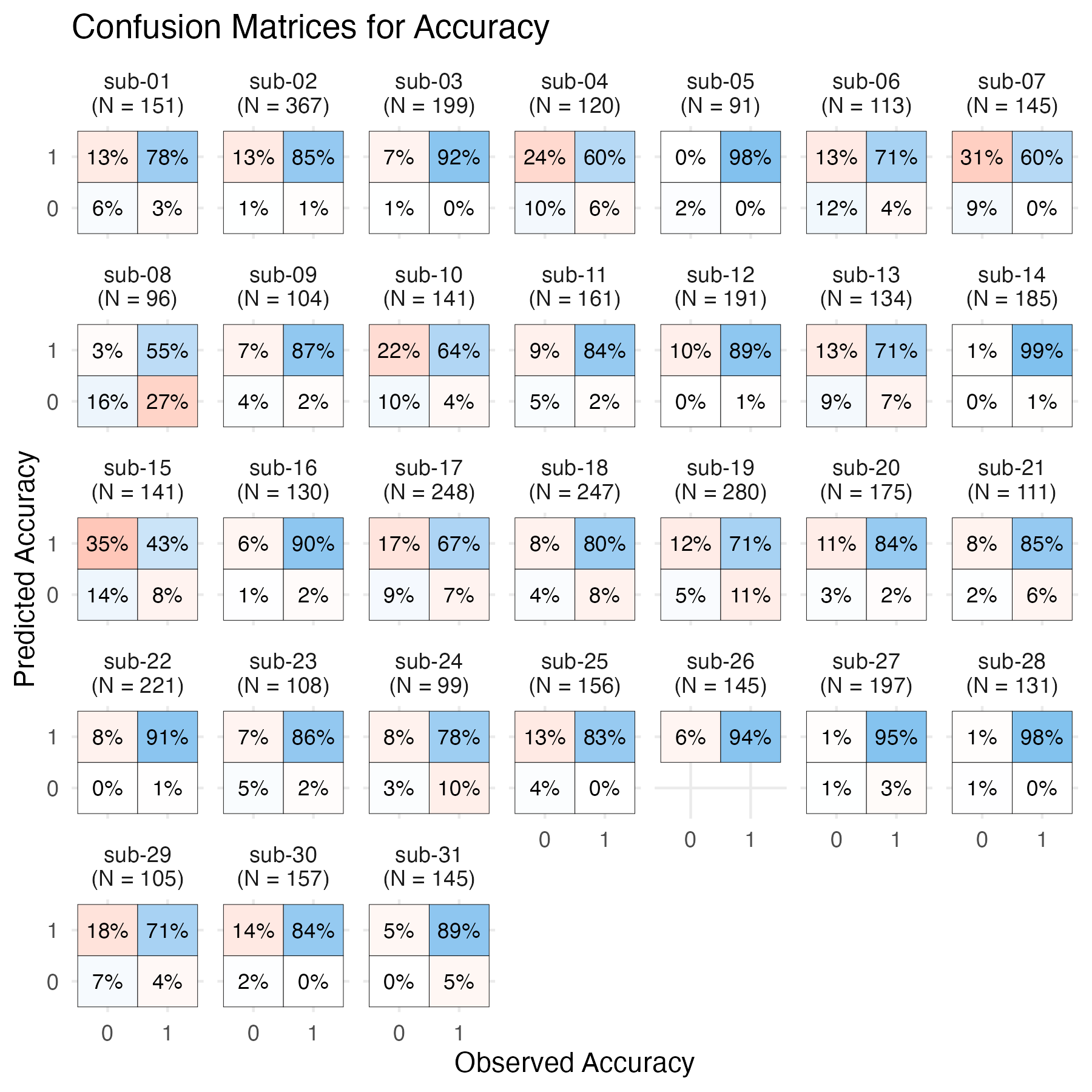

**Figure S1**: *Individual confusion matrices showing the different combinations of predicted and observed accuracies (0 = incorrect, 1 = correct). Numbers indicate the percentage of trials in each cell; colors indicate whether the prediction matched (blue palette) or mismatched (red palette) the observed trial accuracy.*

Because of the task’s adaptive design, participants answered most trials correctly, resulting in a ceiling effect. Thus, the individual confusion matrices of Figure S2 might not be very informative. As a result, we also examined the model’s ability to predict each participant’s mean accuracy rather than single-trial classifications. Figure S2 illustrates the correlation between each participant’s observed and predicted mean accuracy across the entire test. As expected, the model, even at its default parameters, successfully captured individual differences in participant accuracy (*r*(32) = 0.49, *p* = 0.005).

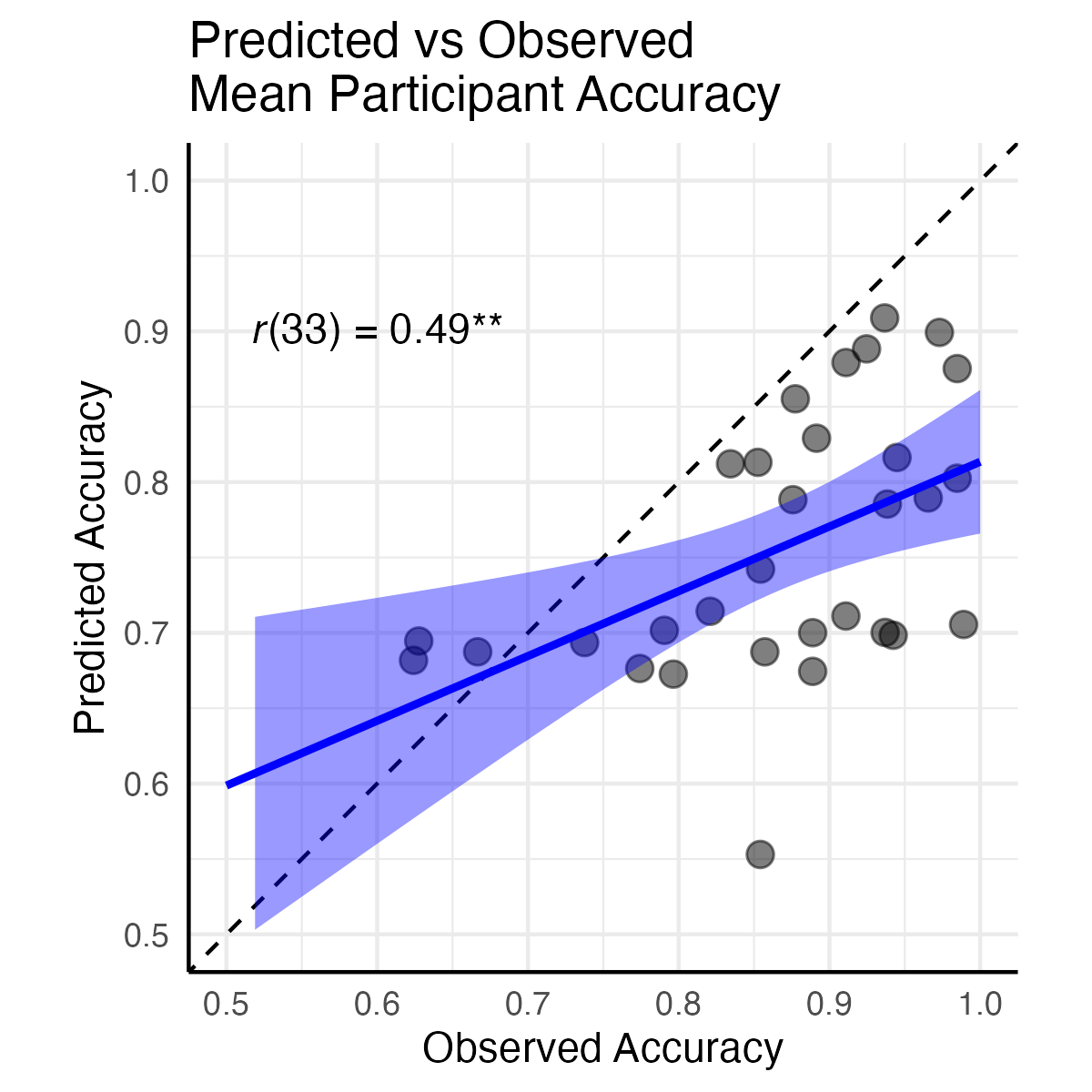

**Figure S2**: *Predicted vs. observed mean accuracies across all trials for each participant. Points represent individual participants; the solid blue line represents the best-fitting regression model; the shaded areas represent 95% confidence intervals of the estimate.*

#### Predicted vs. Observed Response Times

To compare predicted and observed response times, we similarly applied Eq. 4 to the activation associated with the presentation of every test cue, using the default parameters (*t*_0_ = 300 ms, *F* = 1). Figure 6 in the manuscript illustrates the predicted vs. observed distributions of response times, while Figure S3 illustrates the correlations between predicted and observed RTs for every trial for every participant. Individual correlations ranged between 0.19 and 0.62, and were all significant at *p* < 0.05. Across all trials, the mean correlation was *r*(4,844) = 0.36 (*p* < 0.001).

As in the case of accuracy, it is helpful to also look at the model’s capacity to capture variability between, rather than within, individuals. To this end, we computed each participant’s mean response times across all trials and compared each value to the mean of the response times predicted by the model. Figure S4 provides an overview of this analysis; as the figure shows, the model predicts individual variability in mean response times very closely (*r*(33) = 0.84, *p* < 0.001).

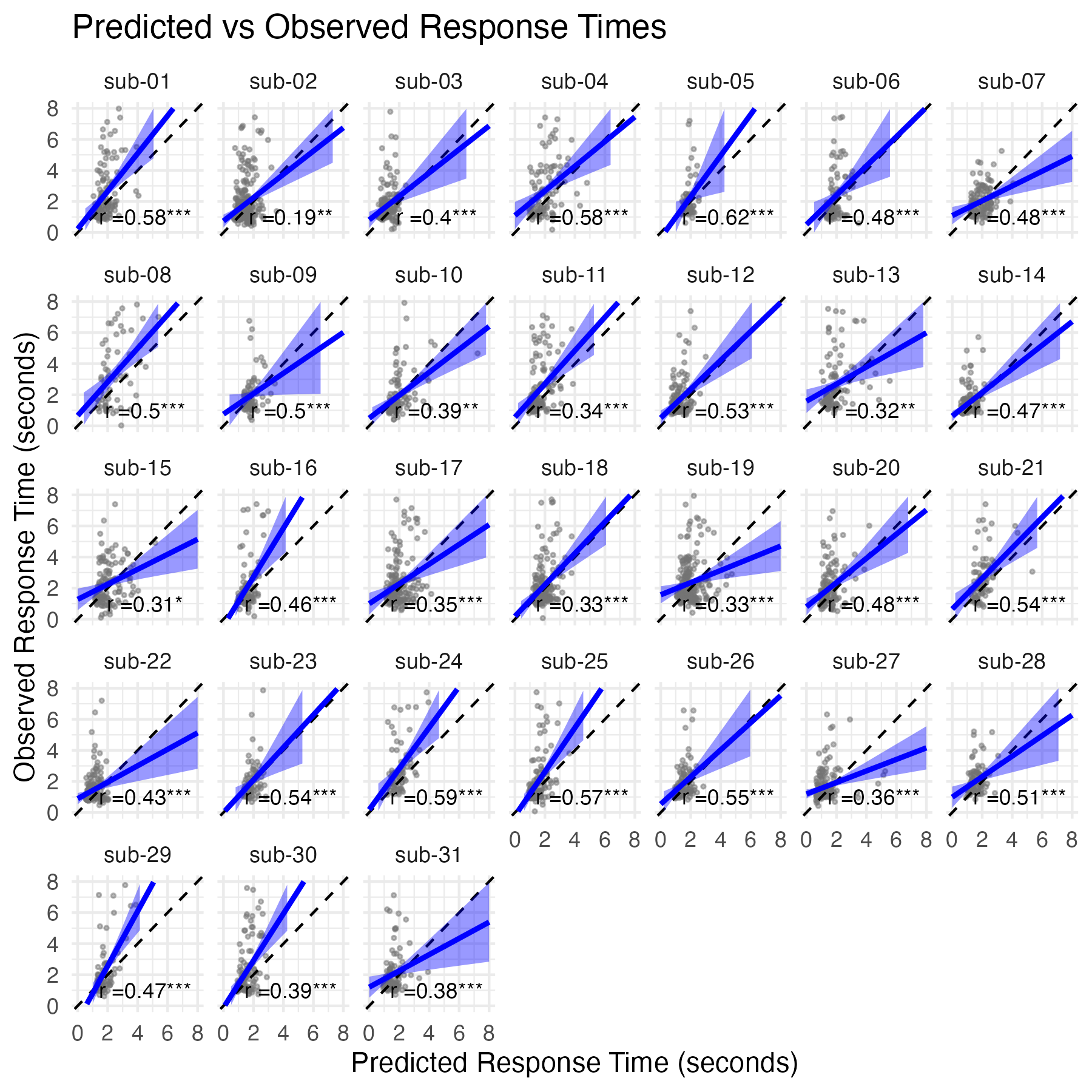

**Figure S3**: *Individual scatterplots showing the predicted and observed responses times for each test trial of each participant, together with their Spearman correlation values (* p < 0.05; ** p < 0.01; *** p < 0.001). Points represent individual trial response times, solid lines represent the best-fitting regression models, shaded areas represent 95% confidence intervals.*

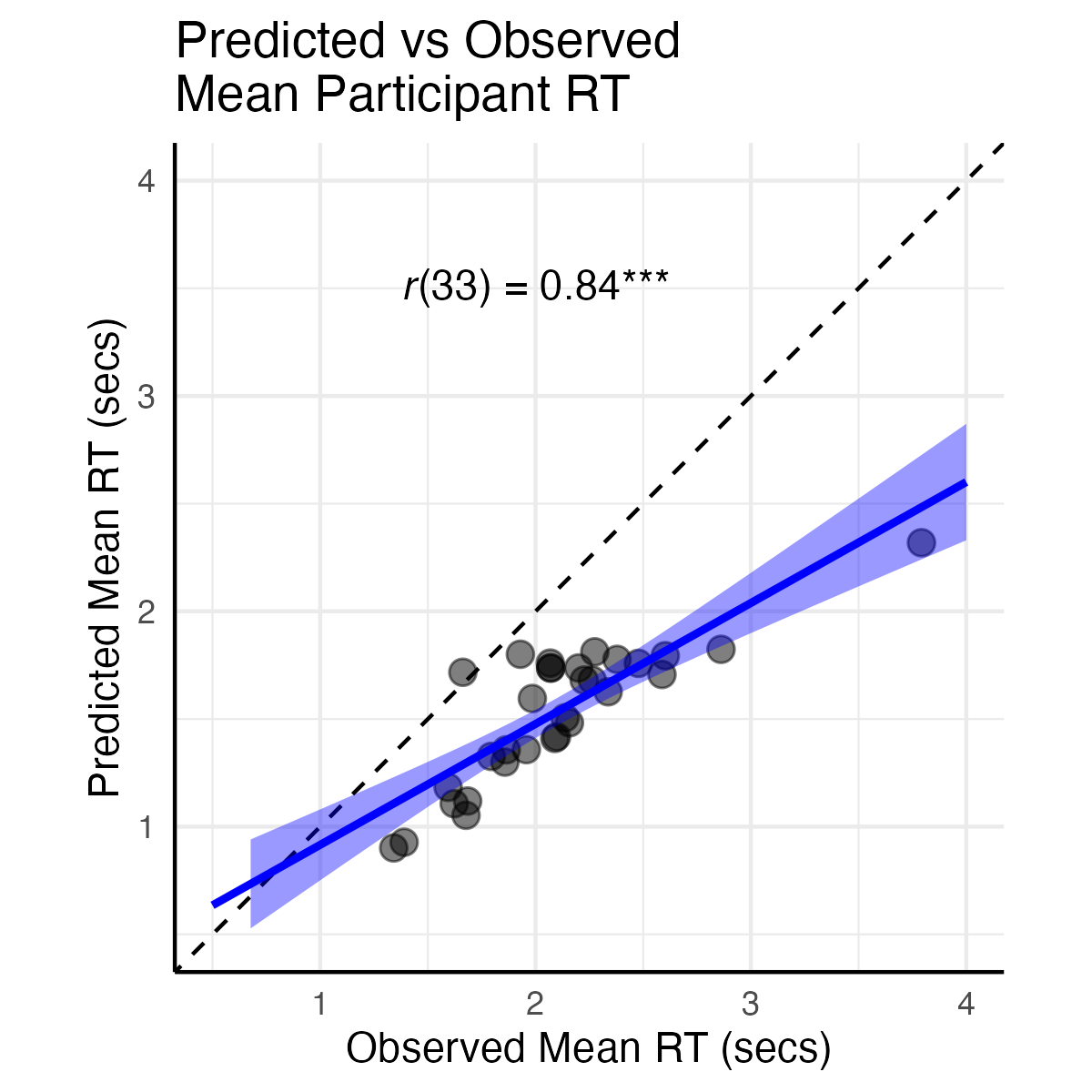

**Figure S4**. *Mean observed responses times for each participant compared the predicted mean response times generated by the model, together with their Spearman correlation value (* p < 0.05; ** p < 0.01; *** p < 0.001). Points represent individual participants, solid lines represent the best-fitting regression models, shaded areas represent 95% confidence intervals.*
